# Use of >100,000 NHLBI Trans-Omics for Precision Medicine (TOPMed) Consortium whole genome sequences improves imputation quality and detection of rare variant associations in admixed African and Hispanic/Latino populations

**DOI:** 10.1101/683201

**Authors:** Madeline H. Kowalski, Huijun Qian, Ziyi Hou, Jonathan D. Rosen, Amanda L. Tapia, Yue Shan, Deepti Jain, Maria Argos, Donna K. Arnett, Christy Avery, Kathleen C. Barnes, Lewis C. Becker, Stephanie A. Bien, Joshua C. Bis, John Blangero, Eric Boerwinkle, Donald W. Bowden, Steve Buyske, Jianwen Cai, Michael H. Cho, Seung Hoan Choi, Hélène Choquet, L Adrienne Cupples, Mary Cushman, Michelle Daya, Paul S. de Vries, Patrick T. Ellinor, Nauder Faraday, Myriam Fornage, Stacey Gabriel, Santhi Ganesh, Misa Graff, Namrata Gupta, Jiang He, Susan R. Heckbert, Bertha Hidalgo, Chani Hodonsky, Marguerite R. Irvin, Andrew D. Johnson, Eric Jorgenson, Robert Kaplan, Sharon LR. Kardia, Tanika N. Kelly, Charles Kooperberg, Jessica A. Lasky-Su, Ruth J.F. Loos, Steven A. Lubitz, Rasika A. Mathias, Caitlin P. McHugh, Courtney Montgomery, Jee-Young Moon, Alanna C. Morrison, Nicholette D. Palmer, Nathan Pankratz, George J. Papanicolaou, Juan M. Peralta, Patricia A. Peyser, Stephen S. Rich, Jerome I. Rotter, Edwin K. Silverman, Jennifer A. Smith, Nicholas L. Smith, Kent D. Taylor, Timothy A. Thornton, Hemant K. Tiwari, Russell P. Tracy, Tao Wang, Scott T. Weiss, Lu Chen Weng, Kerri L. Wiggins, James G. Wilson, Lisa R. Yanek, Sebastian Zöllner, Kari N. North, Paul L. Auer, NHLBI Trans-Omics for Precision Medicine (TOPMed) Consortium, TOPMed Hematology & Hemostasis Working Group, Laura M. Raffield, Alexander P. Reiner, Yun Li

## Abstract

Most genome-wide association and fine-mapping studies to date have been conducted in individuals of European descent, and genetic studies of populations of Hispanic/Latino and African ancestry are still limited. In addition to the limited inclusion of these populations in genetic studies, these populations have more complex linkage disequilibrium structure that may reduce the number of variants associated with a phenotype. In order to better define the genetic architecture of these understudied populations, we leveraged >100,000 phased sequences available from deep-coverage whole genome sequencing through the multi-ethnic NHLBI Trans-Omics for Precision Medicine (TOPMed) program to impute genotypes into admixed African and Hispanic/Latino samples with commercial genome-wide genotyping array data. We demonstrate that using TOPMed sequencing data as the imputation reference panel improves genotype imputation quality in these populations, which subsequently enhances gene-mapping power for complex traits. For rare variants with minor allele frequency (MAF) < 0.5%, we observed a 2.3 to 6.1-fold increase in the number of well-imputed variants, with 11-34% improvement in average imputation quality, compared to the state-of-the-art 1000 Genomes Project Phase 3 and Haplotype Reference Consortium reference panels, respectively. Impressively, even for extremely rare variants with sample minor allele count <10 (including singletons) in the imputation target samples, average information content rescued was >86%. Subsequent association analyses of TOPMed reference panel-imputed genotype data with hematological traits (hemoglobin (HGB), hematocrit (HCT), and white blood cell count (WBC)) in ~20,000 self-identified African descent individuals and ~23,000 self-identified Hispanic/Latino individuals identified associations with two rare variants in the *HBB* gene (rs33930165 with higher WBC (p=8.1×10^−12^) in African populations, rs11549407 with lower HGB (p=1.59×10^−12^) and HCT (p=1.13×10^−9^) in Hispanics/Latinos). By comparison, neither variant would have been genome-wide significant if either 1000 Genomes Project Phase 3 or Haplotype Reference Consortium reference panels had been used for imputation. Our findings highlight the utility of TOPMed imputation reference panel for identification of novel associations between rare variants and complex traits not previously detected in similar sized genome-wide studies of under-represented African and Hispanic/Latino populations.

**Author summary:** Admixed African and Hispanic/Latino populations remain understudied in genome-wide association and fine-mapping studies of complex diseases. These populations have more complex linkage disequilibrium (LD) structure that can impair mapping of variants associated with complex diseases and their risk factors. Genotype imputation represents an approach to improve genome coverage, especially for rare or ancestry-specific variation; however, these understudied populations also have smaller relevant imputation reference panels that need to be expanded to represent their more complex LD patterns. In this study, we leveraged >100,000 phased sequences generated from the multi-ethnic NHLBI TOPMed project to impute in admixed cohorts encompassing ~20,000 individuals of African ancestry (AAs) and ~23,000 Hispanics/Latinos. We demonstrated substantially higher imputation quality for low frequency and rare variants in comparison to the state-of-the-art reference panels (1000 Genomes Project and Haplotype Reference Consortium). Association analyses of ~35 million (AAs) and ~27 million (Hispanics/Latinos) variants passing stringent post-imputation filtering with quantitative hematological traits led to the discovery of associations with two rare variants in the *HBB* gene; one of these variants was replicated in an independent sample, and the other is known to cause anemia in the homozygous state. By comparison, the same *HBB* variants would not have been genome-wide significant using other state-of-the-art reference panels due to lower imputation quality. Our findings demonstrate the power of the TOPMed whole genome sequencing data for imputation and subsequent association analysis in admixed African and Hispanic/Latino populations.

## Introduction

Genotype imputation, despite being a standard practice in modern genetic association studies, remains challenging in populations of Hispanic/Latino or African ancestry, particularly for rare variants (1–6). One obstacle lies in the lack of appropriate whole genome sequence reference panels for these admixed populations. For individuals of European descent, the relevant haplotypes available have increased by more than 500 times from 120 phased sequences in HapMap2 (7) to more than 64,000 phased sequences in Haplotype Reference Consortium (HRC) (8) reference. However, HRC is predominantly European (other than included 1000 Genomes Project Phase 3 (1000G) SNPs) and includes mostly low coverage sequencing data (4-8x coverage). The state-of-the-art reference panels for African ancestry (AA) and Hispanic/Latino cohorts, including the 1000 Genomes Project Phase 3 (1000G) (9) and the Consortium on Asthma among African ancestry Populations in the Americas (CAAPA) (10), are at least one order of magnitude smaller than HRC. This is especially problematic given the complex LD structure in admixed populations. The NHLBI Trans-Omics for Precision Medicine (TOPMed) Project has recently generated deep-coverage (mean depth 30x) whole genome sequencing (WGS) on more than 50,000 individuals from >26 cohorts and from diverse ancestral backgrounds (notably including ~26% AA and ~10% Hispanic/Latino participants), and now provides an unprecedented opportunity for substantially enhancing imputation quality in under-represented admixed populations and subsequently boosting power for mapping genes and regions underlying complex traits. Here we demonstrate the improvements in rare variant imputation quality in AA and Hispanic/Latino populations using TOPMed as a reference panel versus 1000G and HRC panels, and subsequently identify two low frequency/rare *HBB* variant associations with blood cell traits in AA and Hispanic/Latino samples using TOPMed-imputed genotyping array data.

## Results and Discussion

The cohort and ancestry composition of the TOPMed freeze 5b whole genome sequence reference panel used in our study and the samples with array-based genotyping used for imputation and hematological traits association analyses in self-identified AA and Hispanic/Latino individuals are summarized in Tables S1 and S2 respectively. We first selected two large U.S. minority cohorts, one AA and one Hispanic/Latino, in order to comprehensively evaluate imputation quality: the Jackson Heart Study (JHS, all AA, n = 3,082) and the Hispanic Community Health Study/Study of Latinos (HCHS/SOL, all Hispanic/Latino, n = 11,887). Both the JHS and HCHS/SOL have external sources of dense genotype data available for comparison. JHS is the largest AA general population cohort sequenced in TOPMed freeze 5b. Therefore, we removed JHS samples from the TOPMed freeze 5b reference panel prior to performing imputation into JHS samples using SNPs genotyped on the Affymetrix 6.0 array, treating the TOPMed freeze 5b calls as true genotypes for evaluation of imputation quality in JHS. HCHS/SOL is the largest and most regionally diverse population-based cohort of Hispanic/Latino individuals living in the US. For HCHS/SOL, we used the entire set of 100,506 phased sequences from TOPMed freeze 5b (including JHS) as reference, and performed imputation into 11,887 Hispanic/Latino samples genotyped on the Illumina Omni 2.5 SOL custom array (with high quality genotypes at 2,293,536 markers). As the external source of genotype validation in HCHS/SOL, we used genotypes from the Illumina MEGA array genotyping data (containing >1.7 million multi-ethnic global markers, including low frequency coding variants and ancestry-specific variants) available in the same HCHS/SOL samples to assess imputation quality, evaluating 688,189 imputed markers available on MEGA but not on Omni2.5.

Compared with the 1000G Phase 3 reference panelv(9), we were able to increase the number of well-imputed variants from ~28 and ~35 million to ~51 and ~58 million in JHS and HCHS/SOL, respectively (see table S7 for genome-wide distribution of well-imputed variants). We defined well-imputed variants based on our previous work(1, 2, 4), using minor allele frequency (MAF) specific estimated R^2^ thresholds to ensure an average R^2^ of at least 0.8 in each imputed cohort separately. For all rare variants with MAF < 0.5%, we observed ~4.2X (2.3X) and ~6.1X (3.3X) increases in the number of well-imputed variants in JHS (HCHS/SOL), compared with 1000G and HRC respectively, with 22% (11%) and 34% (20%) increases in imputation information content (as measured by average true R^2^, which is the squared Pearson correlation between imputed and true genotypes) (Fig 1 and S1, Table 1). For very rare variants with MAF <0.05%, we observed ~22.1X (5.8X) and ~11.8X (10.7X) increases in the number of well-imputed variants, with 6% (5%) and 13% (11%) increases in average true R^2^, in JHS (HCHS/SOL), compared with 1000G and HRC respectively.

**Table 1.** Number of well-imputed variants using TOPMed freeze 5b, 1000 Genomes Phase 3 (1000G) and Haplotype Reference Consortium (HRC)

| Imputation<br>Reference<br>Panel | Total<br>number of<br>variants in<br>reference<br>panel | Total number of well<br>imputed variants |  | Total number of well imputed variants<br>with MAF<.5% |  |  |  | Total number of well imputed variants<br>with MAF<.05% |  |  |  |
| --- | --- | --- | --- | --- | --- | --- | --- | --- | --- | --- | --- |
|  |  | JHS | HCHS/SOL | JHS | avgTrueR<sup>2</sup> | HCHS/SOL | avgTrueR<sup>2</sup> | JHS | avgTrueR<sup>2</sup> | HCHS/SOL | avgTrueR<sup>2</sup> |
| TOPMed5b | 88,062,238 | 51,467,522 | 57,845,194 | 33,355,468 | 89.89% | 44,439,594 | 89.99% | 16,205,279 | 88.64% | 28,230,718 | 75.21% |
| 1000G* | 49,143,605 | 28,454,330 | 35,178,969 | 7,857,211 | 73.60% | 19,192,645 | 81.08% | 734,063 | 83.72% | 4,901,159 | 71.39% |
| HRC* | 39,635,008 | 21,745,746 | 14,271,587 | 5,488,848 | 67.33% | 13,330,317 | 75.19% | 1,371,526 | 78.77% | 2,637,393 | 67.78% |

HCHS/SOL, Hispanic Community Health Study/Study of Latinos, JHS, Jackson Heart Study, MAF, minor allele frequency
The total number of well imputed variants is extrapolated from three selected 3 Mb regions: 16-19Mb region from chromosomes 3, 12, and 20. These regions were chosen arbitrarily across a range of chromosome sizes, avoiding centromere, telomere, and low-mappability regions. Imputation was carried out using all typed SNPs +/-1Mb (i.e., 15-20Mb) and quality was evaluated in the core 3Mb region. Post imputation quality control was carried out in seven MAF categories separately: <.05%, .05-.2%, .2-.5%, .5-1%, 1-3%, 3-5%, and >5%. In each MAF category, an estimated R<sup>2</sup> threshold was selected to ensure variants above the threshold have an average estimated R<sup>2</sup> of at least 0.8. These variants constitute the well imputed variants

**Fig 1.**
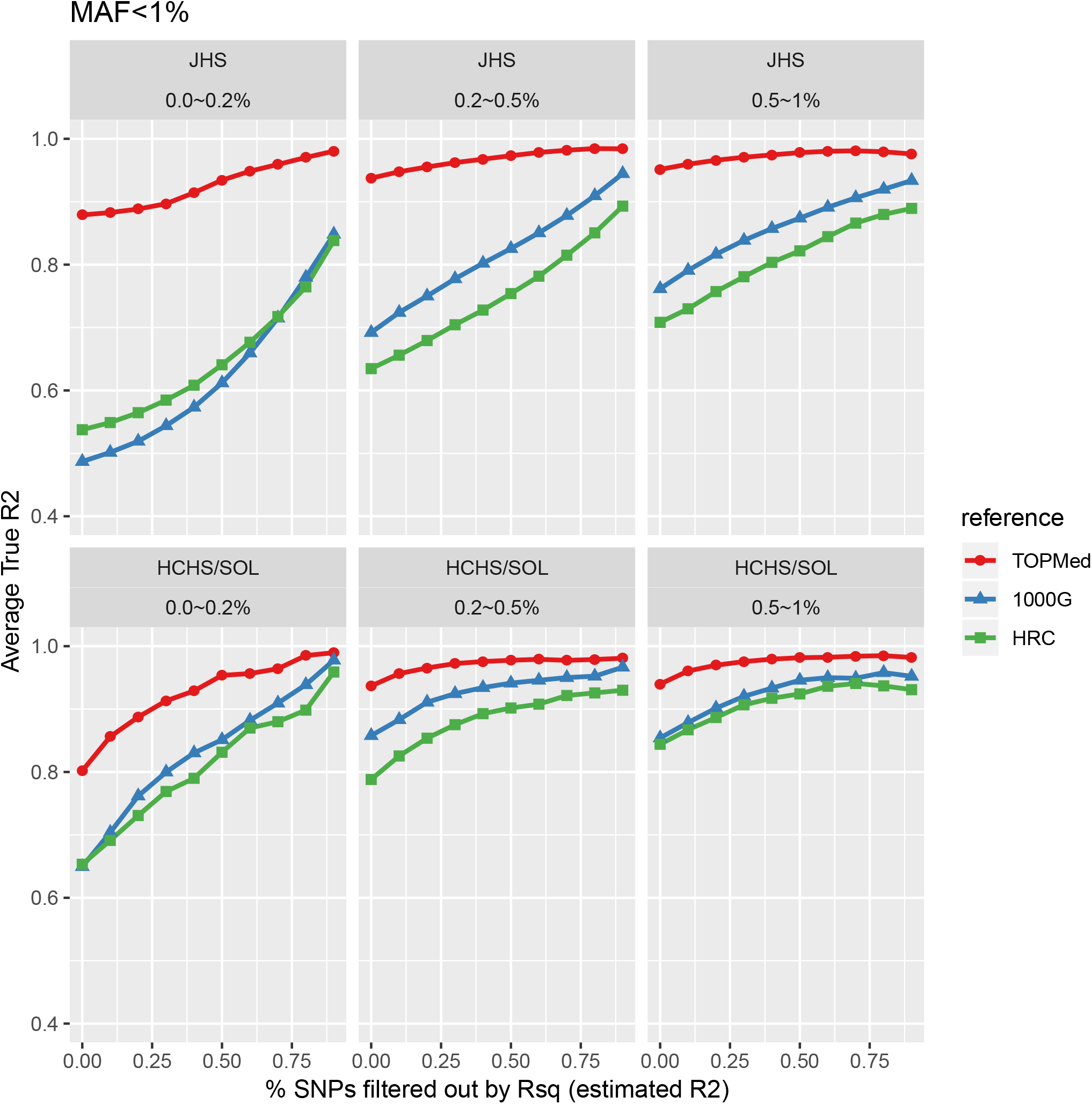
Comparison of imputation reference panels, for variants with MAF < 1%. Imputation quality (measured by true R2 [Y-axis]) is plotted with progressively more stringent post-imputation filtering from left to right, with filtering according to estimated R2 (X-axis), for variants with MAF < 1%. Top panels are for the JHS cohort and bottom panels for the HCHS/SOL cohort. Three reference panels are shown: TOPMed (TOPMed freeze 5b), 1000G (the 1000 Genomes Phase 3), and HRC (the Haplotype Reference Consortium).

Even for extremely rare variants with sample minor allele count (MAC) <10 (including cohort singleton variants in the target JHS cohort), average information content rescued (again measured by true R^2^) was >86%. For example, out of the 8.67 million singleton variants discovered in JHS by TOPMed WGS, 72% (or 6.24 million) can be well-imputed using Affymetrix 6.0 genotypes and using TOPMed freeze 5b (without JHS individuals) as reference, with an average true R^2^ of 0.92 (Table 2). Singletons within JHS are defined as variants with minor allele count of 1 among the JHS samples but which are present in multiple copies in the reference panel. Specifically, average reference MAC is 29.3 before post imputation quality control (QC) and 31.0 after QC, with all variants having a MAC>5 in the overall reference panel. Imputation quality is similarly high when examining extremely rare MAC variants in the reference panel, and even higher as expected with higher MAC variants within the JHS sample (Tables S3-4). Similar observations hold true for HCHS/SOL, with slightly lower imputation quality (Tables S5-6). Compared to JHS African Americans, the lower imputation quality in HCHS/SOL Hispanic/Latino individuals is likely attributable to multiple reasons including (1) the more complex LD structure among Hispanic/Latino individuals due to the presence of three ancestral populations; (2) the availability of a much smaller subset of rare variants for quality evaluation through MEGA array genotyping in HCHS/SOL (in contrast to the availability of nearly all segregating variants in JHS through high coverage sequencing); and (3) the smaller number of relevant haplotypes in the TOPMed freeze 5b reference (~26% self-identified AAs compared to ~10% self-identified Hispanics/Latinos). We note that greater numbers of AA and Hispanic/Latino individuals will be included in future releases of sequencing datasets from TOPMed, which we anticipate will further improve imputation quality; inclusion of JHS itself in imputation for other AA cohorts would also improve imputation quality.

**Table 2.**
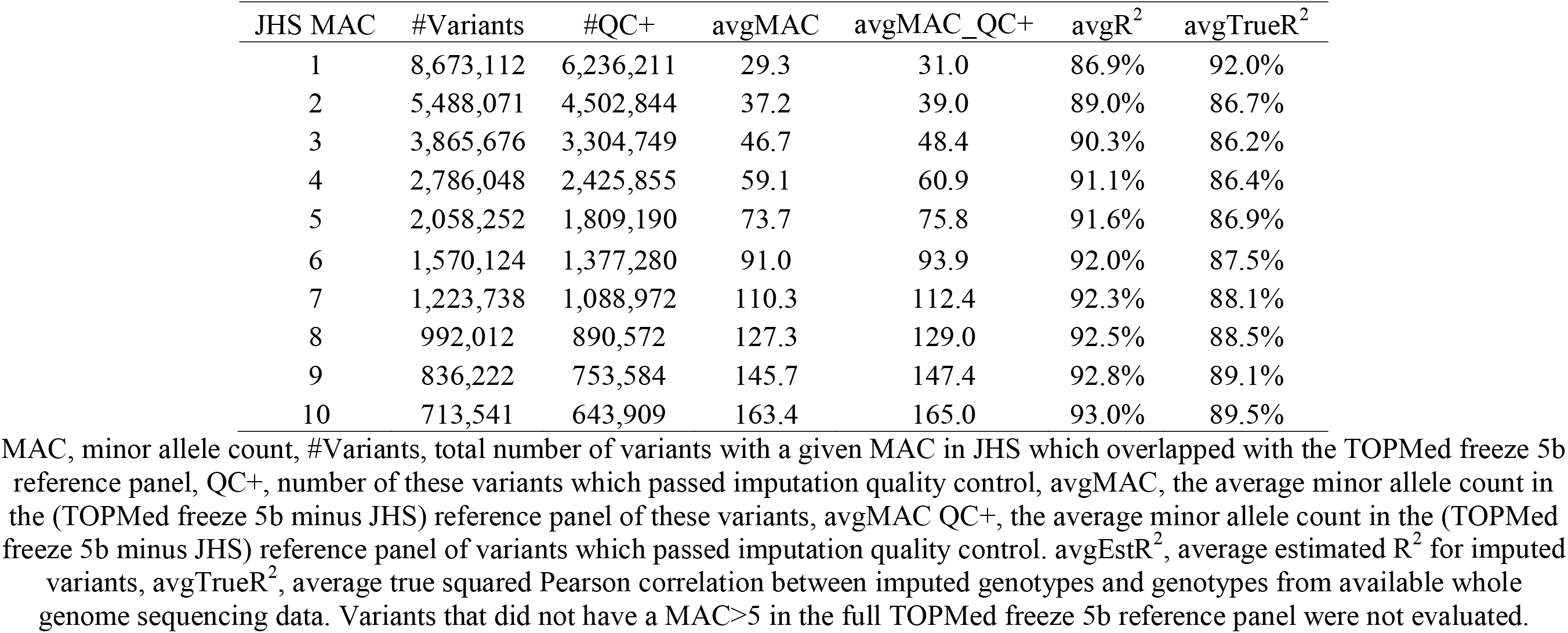
Imputation quality for rare variants (minor allele count<=10) in the Jackson Heart Study (JHS)

Encouraged by these substantial gains in information content for low-frequency and rare variants, we proceeded with imputation in several additional AA and Hispanic/Latino data sets with array-based genotyping (Table S1, S8), followed by association analyses with quantitative blood cell traits to evaluate the power of TOPMed freeze 5b based imputation in minorities for discovery of genetic variants underlying complex human traits. We specifically chose hematological traits for several reasons. First, these traits are important intermediate clinical phenotypes for a variety of cardiovascular, hematologic, oncologic, immunologic, and infectious diseases(11). Second, these traits have family-based heritability estimates in the range of 40-65% (12, 13), and have been highly fruitful for gene-mapping with >2700 common and rare variants identified, though primarily in individuals of European ancestry (14–19). Third, these traits remain under-studied in admixed AA and Hispanic/Latino populations, despite evidence for the existence of variants with distinct genetic architecture in AAs and Hispanics/Latinos (20–22). For example, while hundreds of variants identified in genome-wide association studies (GWAS) of WBC in individuals of European descent explain only ~7% of array heritability, the African specific Duffy null variant DARC rs2814778 alone accounts for 15-20% of population-level WBC variability in AAs(23). Finally, we have previously successfully leveraged deep-coverage exome sequencing-based imputation using resources from the Exome Sequencing Project for more powerful mapping of genes and regions associated with hematological traits in AAs(1). Hemoglobin level (HGB), hematocrit (HCT), and white blood cell count (WBC) were chosen for our primary phenotypic analysis because these traits are available in the largest sample size among the AA and Hispanics/Latinos included in our discovery cohorts.

Our imputation sample used for discovery blood cell trait association analyses included eight cohorts (23,869 AAs and 23,059 Hispanics/Latinos). These discovery samples do not overlap with individuals sequenced as part of TOPMed freeze 5b (Table S2). We used the full set of 100,506 phased sequences from TOPMed freeze 5b (including JHS) as the imputation reference panel. We then carried out AA- and Hispanic/Latino-stratified association analyses with quantitative HGB, HCT, and total WBC separately in each cohort genotyping array data set, accounting for ancestry and relatedness. The genome-wide association results for each imputed cohort data set were then meta-analyzed within each ancestry group. Figs S2-7 show the Manhattan plots from ethnic-specific meta-analyses for each trait. QQ plots (Figs S8-13) show no obvious early departure with genomic control lambda ranging from 0.997 to 1.034, indicating no global inflation of test statistics. For replication of any novel associations identified in the imputation-based discovery analysis, we utilized WGS genotype data and hematological trait data from the non-overlapping set of AA individuals within TOPMed freeze 5b (Table S9) (see Methods for details).

We first evaluated association statistics for variants previously associated with HGB, HCT, or WBC count in AA and Hispanic/Latino populations (summarized in Table S10). We assembled a list of 24 AA and 13 Hispanic/Latino previously identified autosomal signals from prior published GWAS or exome-based studies (1, 19, 20, 24–30). Our lists excluded variants reported in multi-ethnic cohorts or meta-analysis including individuals of non-AA or non-Hispanic/Latino ancestry to guard against the scenario that the reported signals were driven predominantly by individuals of European or Asian ancestry. Among the previously reported 24 AA and 13 Hispanic/Latino variants, all but five (four SNPs and a 3.8 Kb deletion variant esv2676630) passed variant quality control filters in TOPMed freeze 5b and were subsequently well-imputed in our target AA and Hispanic/Latino data sets with a stringent post-imputation R^2^ filter of >0.8 (detailed in Table S11). Among the 32 known HGB, HCT, or WBC count associations testable with TOPMed freeze 5b, our imputed/discovery cohorts confirmed 64.5% of these previously reported findings with a consistent direction of effect, using a stringent genome-wide significant threshold of p<5 × 10^−8^. Using more lenient p-value thresholds, we could replicate 74.2% (p<5 × 10^−6^) and 100% (p<0.05) of the previously reported findings with the same direction of effect. While these results help confirm the overall validity of our hematological trait association results, it is important to note for these comparisons that many of the samples included in the current TOPMed freeze 5b imputed genome-wide association analysis were also used in the publications originally reporting associations in AA and Hispanic/Latino individuals.

Our ancestry-stratified imputation-based discovery meta-analysis revealed two blood cell trait associations that have not been previously reported, at a genome-wide significant threshold of 5×10^-9^ in Hispanics/Latinos and 1×10^−9^ in AA populations, based on appropriate significance thresholds for whole genome sequencing analysis(31). One signal was revealed in each ancestry group: hemoglobin subunit beta (*HBB*) missense (p.Glu7Lys) variant rs33930165 (gb38:11:5227003:C:T) associated with increased WBC in AAs (β=0.31 and p=8.1×10^−12^) (Table 3), and *HBB* stop-gain (p.Gln40Ter) variant rs11549407 (gb38:11:5226774:G:A) associated with lower HGB and HCT in Hispanics/Latinos (β=-1.92 and p=1.59×10^−12^; β=-1.66 and p=1.13×10^−9^). Both variants were either low frequency or rare: the *HBB* missense variant rs33930165 (hemoglobin C variant) has a MAF of 1.14% among the imputed AA discovery samples and is even rarer in non-AA individuals (absent in Europeans in 1000G); the stop gain variant rs11549407 has a MAF of 0.03% (MAC ~ 15) among the imputed Hispanics/Latinos and is monomorphic among the AAs. Both variants are classified as pathogenic in ClinVar. Both variants were well imputed with R^2^ ranging 0.831-0.994 and 0.862-0.999 in the contributing AA and Hispanic/Latino cohorts, respectively (Table 3). Due to the low allele frequency of these variants in AAs and Hispanics/Latinos and even lower frequency in individuals of European descent, both variants were imputed with lower quality using other reference panels (Table S12, Table S13): the missense variant *HBB* rs33930165 had R^2^ as low as 0.127 and 0.456 using 1000G and HRC as reference respectively; the *HBB* stop-gain variant rs11549407 was not available in 1000G and had R^2^ as low as 0.413 using HRC as the reference panel. Carrying the 1000G and HRC imputed genotypes forward to association analyses with hematological traits in the subset of our target imputation cohorts where the variants were well imputed (R^2^ > 0.8), we observed none of the *p-*values exceeded genome-wide significance threshold. This explains why these variants were not detectable at a genome-wide significant level using previously available imputation reference panels.

**Table 3.** Novel variants detected in TOPMed freeze 5b imputed Hispanic/Latino and African ancestry cohorts, in association analyses with white blood cell count, hemoglobin, and hematocrit.

| Ancestry | rs# | Estimated $R^2$<sup>1</sup> | Phenotype | Effect allele | EAF | $\beta$ | SE | P-value | Replication $\beta$ | Replication P-value | Gene | Annotation |
| --- | --- | --- | --- | --- | --- | --- | --- | --- | --- | --- | --- | --- |
| African ancestry | rs33930165 | 0.831-0.994 | WBC | T | 1.14% | 0.31 | 0.05 | 8.1E-12 | 0.27 | 4.6E-04 | *HBB* | missense (p.Glu7Lys) |
| Hispanic/Latino | rs11549407 | 0.862-1.000 | HCT | A | 0.03% | -1.66 | 0.27 | 8.77E-10 |  | NA<sup>4</sup> | *HBB* | stop gain (p.Gln40Ter) |
| Hispanic/Latino | rs11549407 | 0.862-1.000 | HGB | A | 0.03% | -1.92 | 0.27 | 1.53E-12 |  | NA<sup>4</sup> | *HBB* | stop gain (p.Gln40Ter) |

EAF, effect allele frequency, HCT, hematocrit, HGB, hemoglobin, WBC, white blood cell count.
Imputation $R^2$ (estimated $R^2$ ) range reported across all included imputed cohorts.
Association results adjusted for nearby known SNPs whenever applicable.
Association models for rs33930165 were adjusted for SNP rs2814778; removing potential minor allele homozygotes
Association models for rs11549407 were adjusted for SNPs rs334, rs33930165, and rs2213169
rs334 and rs2213169 did not pass variant quality filters in TOPMed freeze 5b and were not included in our main analyses. However, to follow up our novel results in the *HBB* locus, we phased the failed variants in freeze5b and performed targeted imputation using TOPMed freeze 5b calls for rs334 and rs2213169
NA: among TOPMed freeze 5b Hispanic/Latino individuals, MAC =1 so association statistics are not available

Both of our genotype-trait associations involve coding variants of *HBB*, which encodes the beta polypeptide chains in adult hemoglobin. The *HBB* stop gain (p.Gln40Ter) variant 11:5226774:G:A or rs11549407 is the most common cause of beta zero thalassemia in West Mediterranean countries, particularly among the founder population of Sardinia(32, 33), where the variant has a population allele frequency of ~5%. The Sardinian population is represented in the HRC reference panel (~3500 individuals), which likely contributes to the reasonable imputation quality observed using HRC in most but not all cohorts, in contrast to the absence of this variant in 1000G, though imputation quality was clearly improved with the TOPMed freeze 5b reference panel. The p.Gln40Ter mutation is much less prevalent outside of the Western Mediterranean, but has been detected among individuals with beta thalassemia among admixed populations from Central and South America(34, 35), which are geographically and genetically similar to some of the Hispanic/Latino samples included in our imputation-based discovery sample. While the individuals carrying the *HBB* p.Gln40Ter allele in our unselected population-based Hispanic/Latino sample were all imputed heterozygotes (consistent with “thalassemia minor” and generally considered healthy), there is increasing evidence that silent carriers of beta-thalassemia and sickle cell mutations may be at risk for various health-related conditions (36, 37). Due to the relatively small number of Hispanic/Latino individuals with blood cell trait data in TOPMed freeze 5b (n~1,080), including only one heterozygote carrier of rs11549407 in those with blood cell traits measured, we were unable to perform a well-powered replication of the association of rs11549407 with HGB and HCT. Moderate anemia is known to occur in some individuals with thalassemia minor, however, concordant with our results (38).

The association of the *HBB* missense (p.Glu7Lys) variant 11:5227003:C:T or rs33930165 with higher total WBC (β =0.28 and p=8.1×10^−12^) among AA was unexpected; rs33930165 has been associated with red blood cell indices such as mean corpuscular hemoglobin concentration (20) but not with white blood cell traits. Because of the higher allele frequency of this variant and also the larger number of AA samples (n=6,743) in TOPMed freeze 5b, we were able to replicate this *HBB* rs33930165 association with total WBC in an independent sample (β=0.27 and p=4.6×10^−4^) of AA individuals. By contrast, there was no significant association of the *HBB* rs33930165 p.Glu7Lys variant with HGB and a modest association with lower HCT in the AA discovery and replication data sets (discovery HCT β=-0.102, p=0.033, HGB β=0.035, p=0.463, replication HCT β=-0.239, p=0.002, HGB β=-0.009, p=0.909). The minor allele T of rs33930165 encodes an abnormal form of hemoglobin, Hb C, which in the homozygous state is associated with mild chronic hemolytic anemia and mild to moderate splenomegaly (39). In our discovery and replication data sets, there were no individuals homozygous for the Hb C variant, nor any compound heterozygotes for Hb S/C (Hb S is sickling form of hemoglobin and individuals homozygous for Hb S have sickle cell disease), which excludes the possibility that the apparently higher WBC is driven by an “inflammatory response” confined to a small number of individuals clinically affected by sickle cell disease or hemoglobin C disease. We next evaluated the association of *HBB* rs33930165 with circulating number of WBC subtypes, including neutrophils, monocytes, lymphocytes, basophils, and eosinophils. Table S14 shows the results in our AA imputation-based discovery data sets (Table S15), and TOPMed freeze 5b WGS replication samples (Table S16), which suggest that the apparent association of *HBB* rs33930165 with total WBC is mainly driven by an association with higher lymphocyte count, with perhaps a more modest association with higher neutrophil count. Further studies are needed to delineate the putative mechanism of this unexpected association.

Our findings showcase the power of the large, ancestrally diverse TOPMed WGS data set as an imputation reference panel for admixed populations, in terms of both imputation quality and accuracy (especially for rare variants) and subsequent association studies for complex traits. Specifically, we identified two rare variants associated with hematological traits in AA and Hispanic/Latino populations and were able to validate our initial *HBB* association with WBC in an independent replication sample of sequenced individuals. We expect the combination of high-quality imputation and higher depth sequencing datasets in larger cohorts of individuals will provide increased power for rare variant association analyses in diverse populations in the near future.

## Methods

### TOPMed 5b Sequencing and Phasing

The reference panel used for imputation was obtained from deep-coverage whole genome sequences derived from NHLBI’s TOPMed program (www.nhlbiwgs.org), freeze 5b (September 2017). This release included 54,035 non-duplicated, dbGaP released samples, of whom 50,253 have consent to be part of an imputation reference panel. The parent studies that contributed these 50,253 samples are listed in Table S2. Specific to our analyses, freeze 5b includes 3,082 individuals from the Jackson Heart Study, who were removed from the reference panel for our analysis of imputation quality in this particular cohort. Overall, freeze 5b included 54% European ancestry, 26% AA, 10% Hispanic/Latino, 7% Asian, and 3% other ancestry samples. Detailed sequencing methods used in TOPMed are available at https://www.nhlbiwgs.org/topmed-whole-genome-sequencing-project-freeze-5b-phases-1-and-2. In brief, WGS with mean genome coverage ≥30x was completed at six sequencing centers (New York Genome Center, the Broad Institute of MIT and Harvard, the University of Washington Northwest Genomics Center, Illumina Genomic Services, Macrogen Corp., and Baylor Human Genome Sequencing Center). Sequence data files were transferred from sequencing centers to the TOPMed Informatics Research Center (IRC), where reads were aligned to human genome build GRCh38, using a common pipeline, and joint genotype calling was undertaken. Variants were filtered using a machine learning based support vector machine (SVM) approach, using variants present on genotyping arrays as positive controls and variants with many Mendelian inconsistencies as negative controls. After filtering potentially problematic variant sites, freeze 5b contained ~438 million single nucleotide polymorphisms and ~33 million short insertion-deletion variants. For our imputation analyses, we excluded from the reference panel variants with an overall allele count of 5 or less (leaving 88,062,238 variants in our reference panel, Table 1). Additional sample level quality control (such as detection of sex mismatches, pedigree discrepancies, sample swaps, etc.) was undertaken by the TOPMed Data Coordinating Center (DCC).

### Genome-wide genotyping array data sets used for evaluation of imputation quality and/or phenotype association analysis

#### Hispanic Community Health Study/Study of Latinos (HCHS/SOL)

The HCHS/SOL cohort began in 2006 as a prospective study of Hispanic/Latino populations in the U.S. (40–42). From 2008 to 2011, 16,415 adults were recruited from a random sample of households in four communities (the Bronx, Chicago, Miami, and San Diego). Each Field Center recruited >4,000 participants from diverse socioeconomic groups. Most participants self-identified as having Cuban, Dominican, Puerto Rican, Mexican, Central American, or South American heritage. The cohort has been genotyped both using an Illumina Omni2.5M array (plus 150,000 custom SNP, including ancestry-informative markers, Amerindian population specific variants, previously identified GWAS hits, and other candidate polymorphisms for a total of 2,293,715 SNPs)(43) and using the Illumina Multi-Ethnic Genotyping Array (MEGA) array (containing a total of 1,705,969 SNPs) in efforts from the Population Architecture for Genetic Epidemiology(44) consortium to better assess variation in non-European populations. The MEGA array also includes additional exonic, functional, and clinically-relevant variants. Illumina 2.5M array genotypes were available for 12,802 samples, among whom 11,887 samples also had MEGA array genotypes. The Illumina Omni2.5M array was used for imputation to the TOPMed reference panel, with the MEGA array treated as true genotypes for evaluation of imputation quality. For association analysis, imputation was performed on 11,887 samples after merging Omni2.5M array genotypes and MEGA array genotypes (MEGA genotypes were used for variants in both arrays, which resulted in 2,144,214 variants after quality control). For the hematological traits association analysis, 11,588 Hispanic/Latino participants were included.

#### Women’s Health Initiative

The Women’s Health Initiative (WHI) (45) is a long-term national health study focused heart disease, cancer, and osteoporotic fractures in older women. WHI originally enrolled 161,808 women aged 50-79 between 1993 and 1998 at 40 centers across the US, including both a clinical trial (including three trials for hormone therapy, dietary modification, and calcium/vitamin D) and an observational study arm. The recruitment goal of WHI was to include a socio-demographically diverse population with racial/ethnic minority groups proportionate to the total minority population of US women aged 50-79 years. This goal was achieved; a diverse population, including 26,045 (17%) women from minority populations, was recruited. Two WHI extension studies conducted additional follow-up on consenting women from 2005-2010 and 2010-2015. Genotyping was available on some WHI participants through the WHI SNP Health Association Resource (SHARe) resource (dbGaP phs000386.v7.p3), which used the Affymetrix 6.0 array (~906,600 SNPs, 946,000 copy number variation probes) and on other participants through the MEGA array(44). Imputation and association analysis was performed separately in individuals with Affymetrix only, MEGA only, and both Affymetrix and MEGA data (Table S1). For variants with both Affymetrix and MEGA genotypes available, MEGA genotypes were used. In total, 4,318 Hispanic/Latino and 8,494 AA women with blood cell traits were included.

#### UK Biobank

UK Biobank (46) recruited 500,000 people aged between 40-69 years in 2006-2010, establishing a prospective biobank study to understand the risk factors for common diseases such as cancer, heart disease, stroke, diabetes, and dementia). Participants are being followed-up through routine medical and other health-related records from the UK National Health Service. UK Biobank has genotype data on all enrolled participants, as well as extensive baseline questionnaire and physical measures and stored blood and urine samples. Hematological traits were assayed as previously described(14). Genotyping on custom Axiom arrays and subsequent quality control has been previously described(47). Samples were included in our analyses if ancestry self-report was “Black Carribean”, “Black African”,” Black or Black British”, “White and Black Carribean”, “White and Black African”, or “Any Other Black Background”. Variants were selected based on call rate exceeding 95%, HWE p-value exceeding 10^−8^, and MAF exceeding 0.5%. Subsequently, variants in approximate linkage equilibrium were used to generate ten principle components. Samples were excluded if the first principal component exceeded 0.1 and the second principal component exceeded 0.2, to exclude individuals not clustering with most African ancestry individuals. In total, 6,820 AA participants with blood cell traits were included in the analysis.

#### Genetic Epidemiology Research on Aging (GERA)

The GERA cohort includes over 100,000 adults who are members of the Kaiser Permanente Medical Care Plan, Northern California Region (KPNC) and consented to research on the genetic and environmental factors that affect health and disease, linking together clinical data from electronic health records, survey data on demographic and behavioral factors, and environmental data with genetic data. The GERA cohort was formed by including all self-reported racial and ethnic minority participants with saliva samples (19%); the remaining participants were drawn sequentially and randomly from non-Hispanic White participants (81%). Genotyping was completed as previously described(48) using 4 different custom Affymetrix Axiom arrays with ethnic-specific content to increase genomic coverage. Principal components analysis was used to characterize genetic structure in this multi-ethnic sample, as previously described (49). Blood cell traits were extracted from medical records. In individuals with multiple measurements, the first visit with complete white blood cell differential (if any) was used for each participant. Otherwise, the first visit was used. In total, 5,783 Hispanic/Latino and 2,246 AA participants with blood cell traits were included in the analysis.

#### Jackson Heart Study (JHS)

JHS is a population based study designed to investigate risk factors for cardiovascular disease in African Americans. JHS recruited 5,306 AA participants age 35-84 from urban and rural areas of the three counties (Hinds, Madison and Rankin) that comprise the Jackson, Mississippi metropolitan area from 2000-2004, including a nested family cohort (≥ 21 years old) and some prior participants from the Atherosclerosis Risk in Communities (ARIC) study(50, 51). Genotyping was performed using an Affymetrix 6.0 array through NHLBI’s Candidate Gene Association Resource (CARe) consortium(52) in 3,029 individuals, with quality control described previously(53). Due to the greater JHS sample size in TOPMed freeze 5b (n=3,082), we extracted SNPs genotyped on Affymetrix 6.0 and which passed CARe consortium quality control in the non-duplicated JHS TOPMed sequenced samples included in the imputation reference panel (821,172 variants which passed TOPMed quality controls used for imputation).

#### Coronary Artery Risk Development in Young Adults (CARDIA)

The CARDIA study is a longitudinal study of cardiovascular disease risk initiated in 1985-86 in 5,115 AA and European ancestry men and women, then aged 18-30 years. The CARDIA sample was recruited at four sites: Birmingham, AL, Chicago, IL, Minneapolis, MN, and Oakland, CA (54, 55). Similar to JHS, genotyping was performed through the CARe consortium(52, 53) using an Affymetrix 6.0 array. In total, 1,619 AA participants with blood cell traits were included in the analysis.

#### Atherosclerosis Risk in Communities (ARIC)

The ARIC study was initiated in 1987, when participants were 45-64 years old, recruiting participants age 45-64 years from 4 field centers (Forsyth County, NC; Jackson, MS; northwestern suburbs of Minneapolis, MN; Washington County, MD) in order to study cardiovascular disease and its risk factors(56), including the participants of self-reported AA ancestry included here. Standardized physical examinations and interviewer-administered questionnaires were conducted at baseline (1987–89), three triennial follow-up examinations, a fifth examination in 2011-13, and a sixth exam in 2016-2017. Genotyping was performed through the CARe consortium Affymetrix 6.0 array.(52, 53) In total, 2,392 AA participants with blood cell traits were included in the analysis.

### Imputation and post-imputation quality filtering

We first phased individuals from each cohort separately using *eagle(57)* with default settings. We subsequently performed haplotype-based imputation using *minimac4(58)* using phased haplotypes from TOPMed freeze 5b as reference. We used 100,506 TOPMed freeze 5b whole genome sequences as reference for all cohorts except JHS, for which we used 94,342 TOPMed freeze 5b non-JHS sequences. We additionally imputed HCHS/SOL and JHS using 1000 Genomes Phase 3(9) and HRC(8) reference panels. Post-imputation quality filtering was performed using a R^2^ threshold specific to each MAF category to ensure average R^2^ for variants passing threshold was at least 0.8, following our previous work(4, 59). Restricting to variants passing post-imputation quality control in at least two cohorts resulted in 34.4-35.8 million variants assessed in the AA cohorts and 26.7-27.2 million assessed in the HA cohorts, depending on the exact sample size of the tested trait. Imputation and association analysis included autosomal variants only. We assessed imputation quality (comparing true and estimated average R^2^) in three selected 3Mb regions: 16-19Mb region (relative to the start of each chromosome) from chromosomes 3, 12, and 20.

### Hematological traits

HGB, HCT, WBC and differential were measured in both the discovery data sets (Tables S7, S13) and a subset of the TOPMed freeze 5b samples (Tables S8, S14) using automated clinical hematology analyzers. Prior to association analyses, we excluded extreme outlier values, notably WBC values >200×10^9^/L (as well as WBC subtype count values in these individuals), HCT >60%, and HGB >20g/dL. For longitudinal cohort studies, all values are from the same exam cycle, chosen based on largest available sample size. WBC traits were log transformed due to their skewed distribution. For all traits, we first derived trait residuals adjusting for age, age squared, sex, and principal components/study specific covariates as needed. Trait residuals were then inverse-normalized prior to analysis.

### Association analysis in discovery cohorts

Association analyses were carried out for these variants via EPACTS for all cohorts except for HCHS/SOL, using the *q.emmax* test to account for relatedness within each cohort. Association tests were performed on inverse normalized residuals (adjusted for age, age squared, sex, and principal components/study specific covariates), further adjusting for kinship matrices constructed in EPACTS using variants with a MAF>1%. Individuals with different starting genotyping platform(s) were also analyzed separately. Inverse-variance weighted meta-analysis were further carried out using GWAMA(60), separately for AAs and Hispanics/Latinos.

### Identification and replication of novel associations

To identify putative novel associations, we then filtered out any variant with LD r^2^ ≥ 0.2 in any ethnic group with any previous reported variant from GWAS, sequencing, or Exome Chip analyses within ±1Mb for a given blood cell trait. We calculated LD in self-reported European ancestry, AA, and Hispanic/Latino individuals from TOPMed freeze 5b. For European and African LD reference panels, we further restricted to individuals with global ancestry estimate ≥0.8. The global ancestry estimates were derived from local ancestry estimates from RFMix(61) using data from the Human Genome Diversity Project (HGDP)(62) as the reference panel with seven populations, namely Sub-Saharan Africa, Central and South Asia, East Asia, Europe, Native America, Oceania, and West Asia and North Africa (Middle East). Global ancestry for each TOPMed individual is defined as the mean local ancestry across all HGDP SNPs. For replication of novel signals, similar to the approach we adopted for the discovery cohorts, we performed association analysis using EPACTS in each contributing cohort and then meta-analyzed with GWAMA.

## Author Contributions

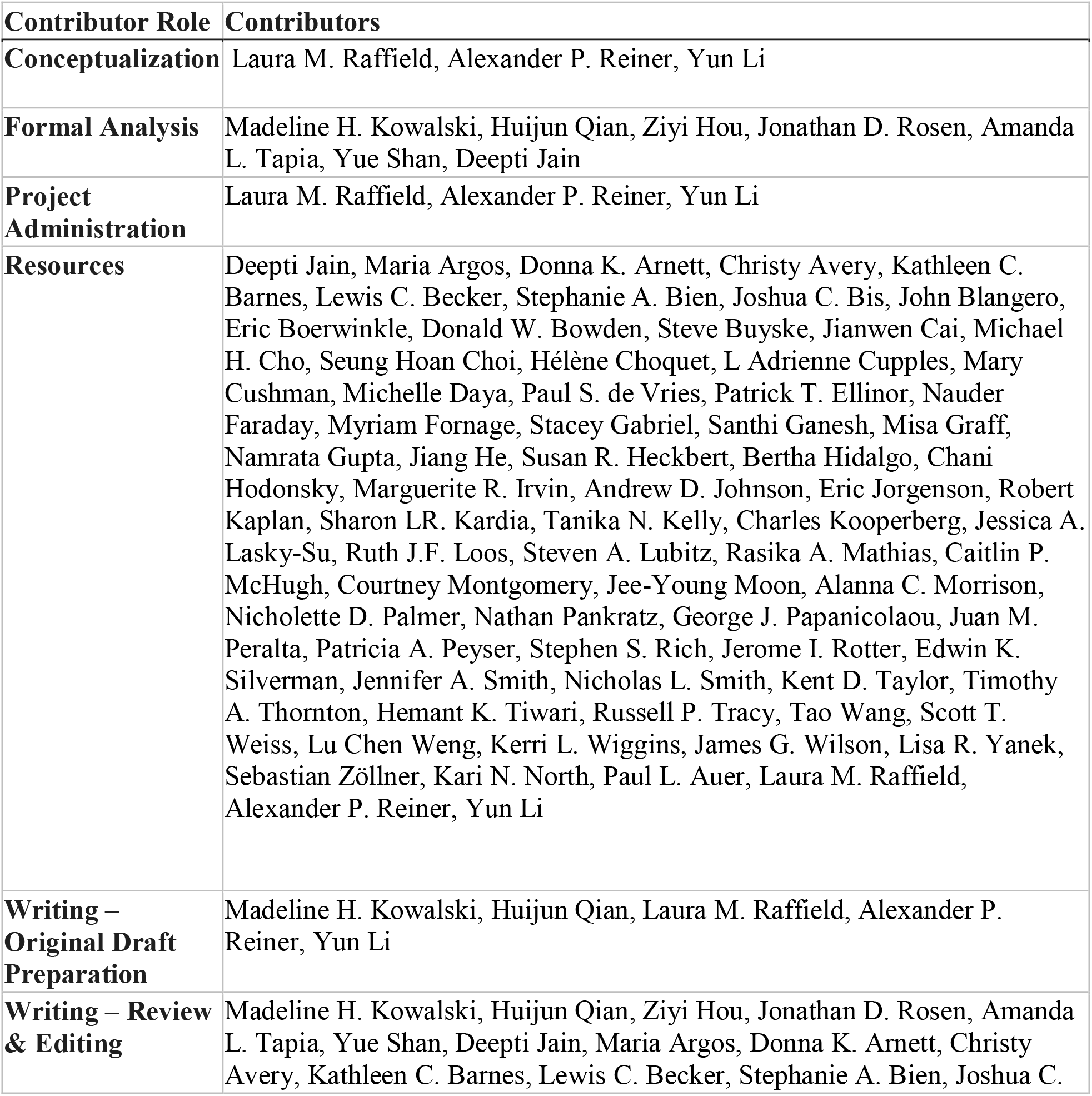
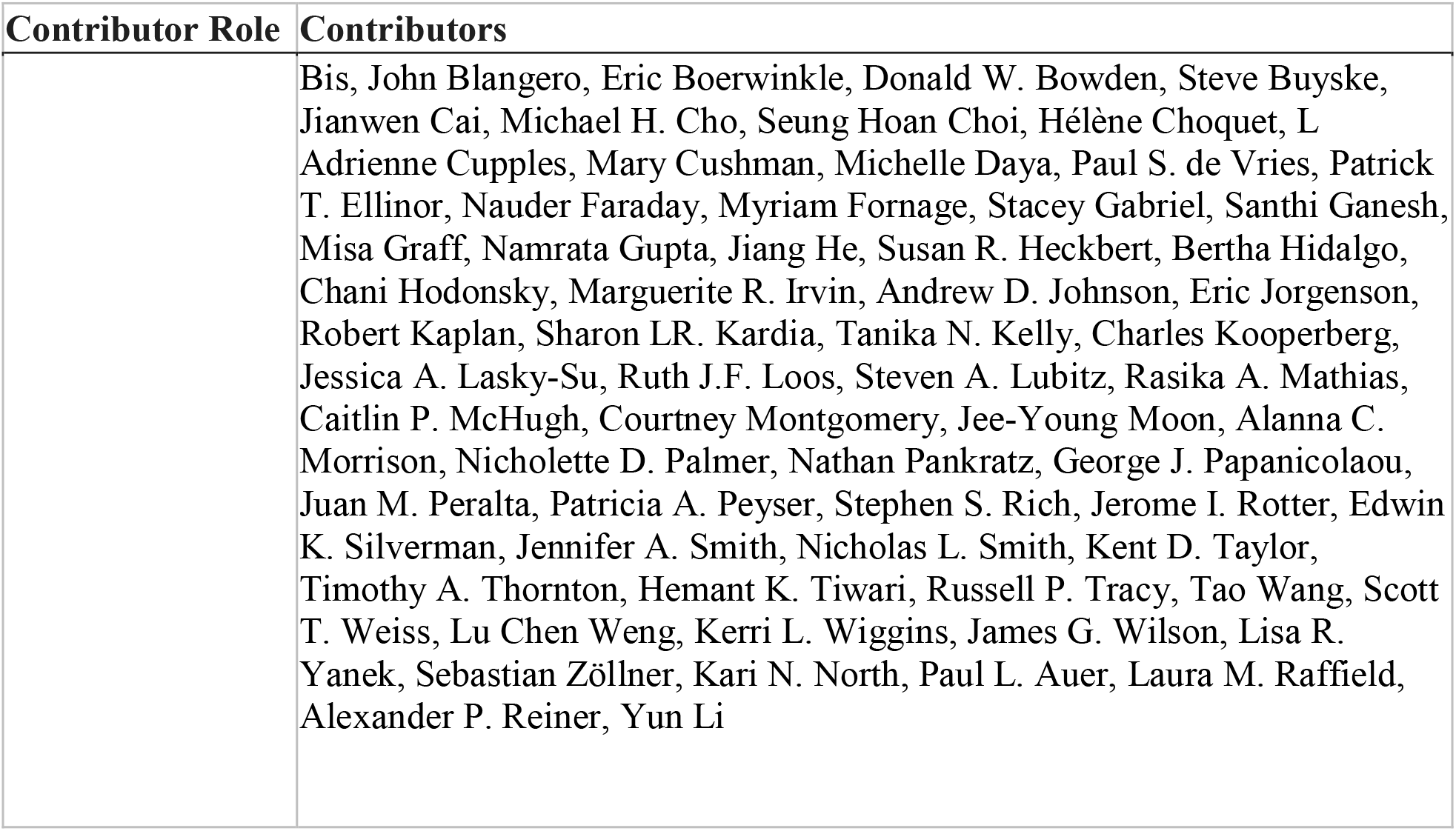

## Acknowledgments

This research has been conducted using the UK Biobank Resource under Application Number 25953.

Support for title page creation and format was provided by AuthorArranger, a tool developed at the National Cancer Institute.

Whole genome sequencing (WGS) for the Trans-Omics in Precision Medicine (TOPMed) program was supported by the National Heart, Lung and Blood Institute (NHLBI). WGS for “NHLBI TOPMed: The Jackson Heart Study” (phs000964) was performed at the University of Washington Northwest Genomics Center (HHSN268201100037C). WGS for “NHLBI TOPMed: Genetic Epidemiology of COPD (COPDGene) in the TOPMed Program” (phs000951) was performed at the Broad Institute and the University of Washington Northwest Genomics Center (HHSN268201500014C (Phase 2 Broad)). WGS for “NHLBI TOPMed: Genetics of Cardiometabolic Health in the Amish” (phs000956) was performed at the Broad Institute (3R01HL121007-01S1). WGS for “NHLBI TOPMed: Whole Genome Sequencing and Related Phenotypes in the Framingham Heart Study” (phs000974) was performed at the Broad Institute (3R01HL092577-06S1 (AFGen)). WGS for “NHLBI TOPMed: The Genetic Epidemiology of Asthma in Costa Rica” (phs000988) was performed at the University of Washington Northwest Genomics Center (3R37HL066289-13S1). WGS for “NHLBI TOPMed: Heart and Vascular Health Study (HVH)” (phs000993) was performed at the Broad Institute and Baylor (3R01HL092577-06S1 (Phase 1:Broad, AFGen), 3U54HG003273-12S2 (Phase 2: Baylor, VTE, TOPMed supplement to NHGRI)). WGS for “NHLBI TOPMed: The Vanderbilt AF Ablation Registry” (phs000997) was performed at the Broad Institute (3R01HL092577-06S1). WGS for “NHLBI TOPMed: Partners HealthCare Biobank” (phs001024) was performed at the Broad Institute (3R01HL092577-06S1). WGS for “NHLBI TOPMed: The Vanderbilt Atrial Fibrillation Registry” (phs001032) was performed at the Broad Institute (3R01HL092577-06S1). WGS for “NHLBI TOPMed: Novel Risk Factors for the Development of Atrial Fibrillation in Women” (phs001040) was performed at the Broad Institute (3R01HL092577-06S1). WGS for “NHLBI TOPMed: MGH Atrial Fibrillation Study” (phs001062) was performed at the Broad Institute (3R01HL092577-06S1). WGS for “NHLBI TOPMed: The Genetics and Epidemiology of Asthma in Barbados” (phs001143) was performed at Illumina (3R01HL104608-04S1). WGS for “NHLBI TOPMed: Cleveland Clinic Atrial Fibrillation Study” (phs001189) was performed at the Broad Institute (3R01HL092577-06S1). WGS for “NHLBI TOPMed: African American Sarcoidosis Genetics Resource” (phs001207) was performed at Baylor (3R01HL113326-04S1). WGS for “NHLBI TOPMed: Trans-Omics for Precision Medicine Whole Genome Sequencing Project: ARIC” (phs001211) was performed at the Broad Institute and Baylor (3R01HL092577-06S1 (Broad, AFGen), HHSN268201500015C (Baylor, VTE), 3U54HG003273-12S2 (Baylor, VTE)). WGS for “NHLBI TOPMed: San Antonio Family Heart Study (WGS)” (phs001215) was performed at Illumina (3R01HL113323-03S1). WGS for “NHLBI TOPMed: Genetic Epidemiology Network of Salt Sensitivity (GenSalt)” (phs001217) was performed at Baylor (HHSN268201500015C). WGS for “NHLBI TOPMed: GeneSTAR (Genetic Study of Atherosclerosis Risk)” (phs001218) was performed at the Broad Institute and Macrogen (HHSN268201500014C (Broad, AA_CAC)). WGS for “NHLBI TOPMed: Women’s Health Initiative (WHI)” (phs001237) was performed at the Broad Institute (HHSN268201500014C). WGS for “NHLBI TOPMed: HyperGEN - Genetics of Left Ventricular (LV) Hypertrophy” (phs001293) was performed at the University of Washington Northwest Genomics Center (3R01HL055673-18S1). WGS for “NHLBI TOPMed: Genetic Epidemiology Network of Arteriopathy (GENOA)” (phs001345) was performed at the Broad Institute and the University of Washington Northwest Genomics Center (HHSN268201500014C (Broad, AA_CAC), 3R01HL055673-18S1 (UW NWGC, HyperGEN_GENOA)). WGS for “NHLBI TOPMed: Genetics of Lipid Lowering Drugs and Diet Network (GOLDN)” (phs001359) was performed at the University of Washington Northwest Genomics Center (3R01HL104135-04S1). WGS for “NHLBI TOPMed: Cardiovascular Health Study” (phs001368) was performed at Baylor (HHSN268201500015C (VTE portion of CHS)). WGS for “NHLBI TOPMed: Whole Genome Sequencing of Venous Thromboembolism (WGS of VTE)” (phs001402) was performed at Baylor (HHSN268201500015C, 3U54HG003273-12S2).WGS for “NHLBI TOPMed: Diabetes Heart Study African American Coronary Artery Calcification (AA CAC)” (phs001412) was performed at the Broad Institute (HHSN268201500014C (Broad, AA_CAC)). WGS for “NHLBI TOPMed: Multi-Ethnic Study of Atherosclerosis (MESA)” (phs001416.v1.p1) was performed at the Broad Institute of MIT and Harvard (3U54HG003067-13S1).WGS for “NHLBI TOPMed: MESA Family AA-CAC” (phs001416) was performed at the Broad Institute (HHSN268201500014C). Centralized read mapping and genotype calling, along with variant quality metrics and filtering were provided by the TOPMed Informatics Research Center (3R01HL-117626-02S1). Phenotype harmonization, data management, sample-identity QC, and general study coordination, were provided by the TOPMed Data Coordinating Center (3R01HL-120393-02S1). We gratefully acknowledge the studies and participants who provided biological samples and data for TOPMed.

The Jackson Heart Study (JHS) is supported and conducted in collaboration with Jackson State University (HHSN268201800013I), Tougaloo College (HHSN268201800014I), the Mississippi State Department of Health (HHSN268201800015I/HHSN26800001) and the University of Mississippi Medical Center (HHSN268201800010I, HHSN268201800011I and HHSN268201800012I) contracts from the National Heart, Lung, and Blood Institute (NHLBI) and the National Institute for Minority Health and Health Disparities (NIMHD). The authors also wish to thank the staffs and participants of the JHS.

MESA and the MESA SHARe project are conducted and supported by the National Heart, Lung, and Blood Institute (NHLBI) in collaboration with MESA investigators. Support for MESA is provided by contracts HHSN268201500003I, N01-HC-95159, N01-HC-95160, N01-HC-95161, N01-HC-95162, N01-HC-95163, N01-HC-95164, N01-HC-95165, N01-HC-95166, N01-HC-95167, N01-HC-95168, N01-HC-95169, UL1-TR-000040, UL1-TR-001079, UL1-TR-001420, UL1-TR-001881, and DK063491. MESA Family is conducted and supported by the National Heart, Lung, and Blood Institute (NHLBI) in collaboration with MESA investigators. Support is provided by grants and contracts R01HL071051, R01HL071205, R01HL071250, R01HL071251, R01HL071258, R01HL071259, and by the National Center for Research Resources, Grant UL1RR033176. The provision of genotyping data was supported in part by the National Center for Advancing Translational Sciences, CTSI grant UL1TR001881, and the National Institute of Diabetes and Digestive and Kidney Disease Diabetes Research Center (DRC) grant DK063491 to the Southern California Diabetes Endocrinology Research Center.

The COPDGene project described was supported by Award Number U01 HL089897 and Award Number U01 HL089856 from the National Heart, Lung, and Blood Institute. The content is solely the responsibility of the authors and does not necessarily represent the official views of the National Heart, Lung, and Blood Institute or the National Institutes of Health. The COPDGene project is also supported by the COPD Foundation through contributions made to an Industry Advisory Board comprised of AstraZeneca, Boehringer Ingelheim, GlaxoSmithKline, Novartis, Pfizer, Siemens and Sunovion. A full listing of COPDGene investigators can be found at: http://www.copdgene.org/directory

GeneSTAR was supported by the National Institutes of Health/National Heart, Lung, and Blood Institute (U01 HL72518, HL087698, HL112064, HL11006, HL118356) and by a grant from the National Institutes of Health/National Center for Research Resources (M01-RR000052) to the Johns Hopkins General Clinical Research Center.

The views expressed in this manuscript are those of the authors and do not necessarily represent the views of the National Heart, Lung, and Blood Institute; the National Institutes of Health; or the U.S. Department of Health and Human Services.

The Population Architecture Using Genomics and Epidemiology (PAGE) program is funded by the National Human Genome Research Institute (NHGRI) with co-funding from the National Institute on Minority Health and Health Disparities (NIMHD). The contents of this paper are solely the responsibility of the authors and do not necessarily represent the official views of the NIH. The PAGE consortium thanks the staff and participants of all PAGE studies for their important contributions. We thank Rasheeda Williams and Margaret Ginoza for providing assistance with program coordination. The complete list of PAGE members can be found at http://www.pagestudy.org.

Assistance with data management, data integration, data dissemination, genotype imputation, ancestry deconvolution, population genetics, analysis pipelines, and general study coordination was provided by the PAGE Coordinating Center (NIH U01HG007419). Genotyping services were provided by the Center for Inherited Disease Research (CIDR). CIDR is fully funded through a federal contract from the National Institutes of Health to The Johns Hopkins University, contract number HHSN268201200008I. Genotype data quality control and quality assurance services were provided by the Genetic Analysis Center in the Biostatistics Department of the University of Washington, through support provided by the CIDR contract. The data and materials included in this report result from collaboration between the following studies and organizations:

**HCHS/SOL:** Primary funding support to Dr. North and colleagues is provided by U01HG007416. Additional support was provided via R01DK101855 and 15GRNT25880008. The Hispanic Community Health Study/Study of Latinos was carried out as a collaborative study supported by contracts from the National Heart, Lung, and Blood Institute (NHLBI) to the University of North Carolina (N01-HC65233), University of Miami (N01-HC65234), Albert Einstein College of Medicine (N01-HC65235), Northwestern University (N01-HC65236), and San Diego State University (N01-HC65237). The following Institutes/Centers/Offices contribute to the HCHS/SOL through a transfer of funds to the NHLBI: National Institute on Minority Health and Health Disparities, National Institute on Deafness and Other Communication Disorders, National Institute of Dental and Craniofacial Research, National Institute of Diabetes and Digestive and Kidney Diseases, National Institute of Neurological Disorders and Stroke, NIH Institution-Office of Dietary Supplements

**WHI:** Funding support for the “Exonic variants and their relation to complex traits in minorities of the WHI” study is provided through the NHGRI PAGE program (NIH U01HG007376). The WHI program is funded by the National Heart, Lung, and Blood Institute, National Institutes of Health, U.S. Department of Health and Human Services through contracts HHSN268201100046C, HHSN268201100001C, HHSN268201100002C, HHSN268201100003C, HHSN268201100004C, and HHSN271201100004C. The authors thank the WHI investigators and staff for their dedication, and the study participants for making the program possible. A listing of WHI investigators can be found at: https://www.whi.org/researchers/Documents%20%20Write%20a%20Paper/WHI%20Investigator%20Short%20List.pdf

**ARIC:** The Atherosclerosis Risk in Communities study has been funded in whole or in part with Federal funds from the National Heart, Lung, and Blood Institute, National Institutes of Health, Department of Health and Human Services (contract numbers HHSN268201700001I, HHSN268201700002I, HHSN268201700003I, HHSN268201700004I and HHSN268201700005I). The authors thank the staff and participants of the ARIC study for their important contributions.

**CARDIA:** The Coronary Artery Risk Development in Young Adults Study (CARDIA) is conducted and supported by the National Heart, Lung, and Blood Institute (NHLBI) in collaboration with the University of Alabama at Birmingham (HHSN268201300025C & HHSN268201300026C), Northwestern University (HHSN268201300027C), University of Minnesota (HHSN268201300028C), Kaiser Foundation Research Institute (HHSN268201300029C), and Johns Hopkins University School of Medicine (HHSN268200900041C). CARDIA is also partially supported by the Intramural Research Program of the National Institute on Aging (NIA) and an intra-agency agreement between NIA and NHLBI (AG0005). This manuscript has been reviewed by CARDIA for scientific content.

**GERA:** Genotyping of the GERA cohort was funded by a grant from the National Institute on Aging, National Institute of Mental Health, and National Institute of Health Common Fund (RC2 AG036607).

## Supporting information

**Fig S1.**
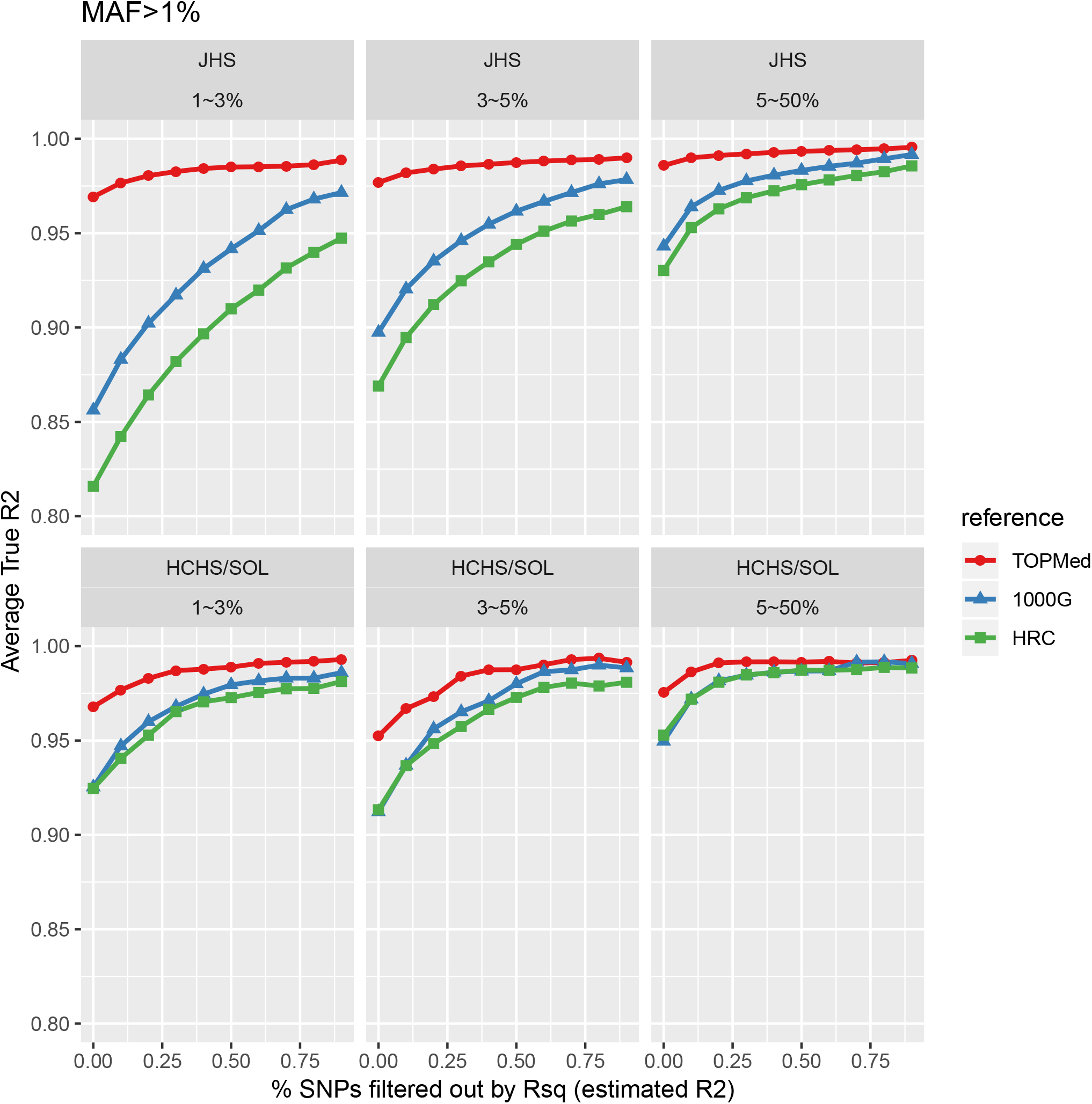
Comparison of imputation reference panels, for variants with MAF > 1%. Imputation quality (measured by true R2 [Y-axis]) is plotted with progressively more stringent post-imputation filtering from left to right, with filtering according to estimated R2 (X-axis), for variants with MAF > 1%. Top panels are for the JHS cohort and bottom panels for the HCHS/SOL cohort. Three reference panels are shown: TOPMed (TOPMed freeze 5b), 1000G (the 1000 Genomes Phase 3), and HRC (the Haplotype Reference Consortium).

**Fig S2.**
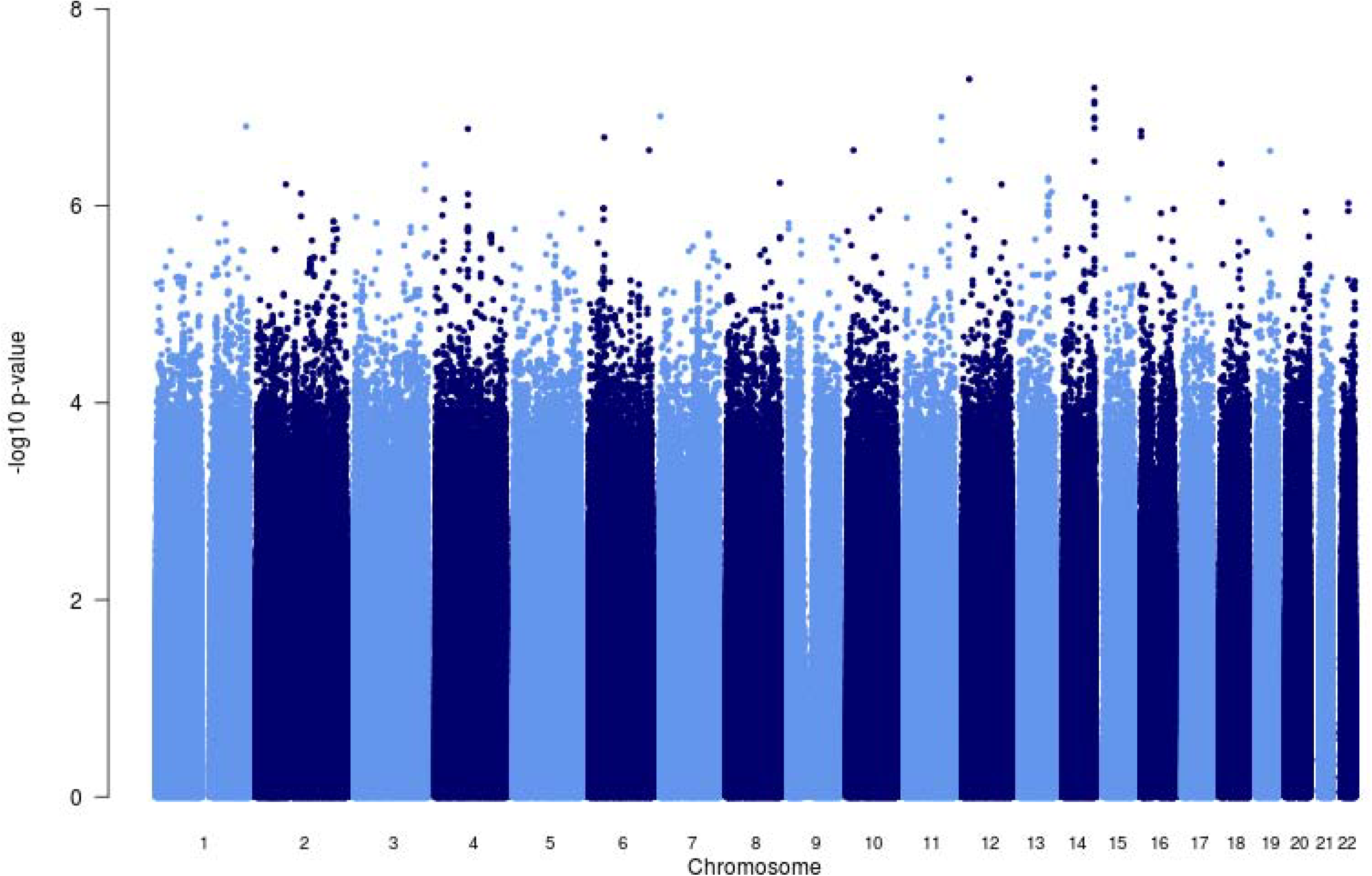
African ancestry hematocrit analysis Manhattan plot.

**Fig S3.**
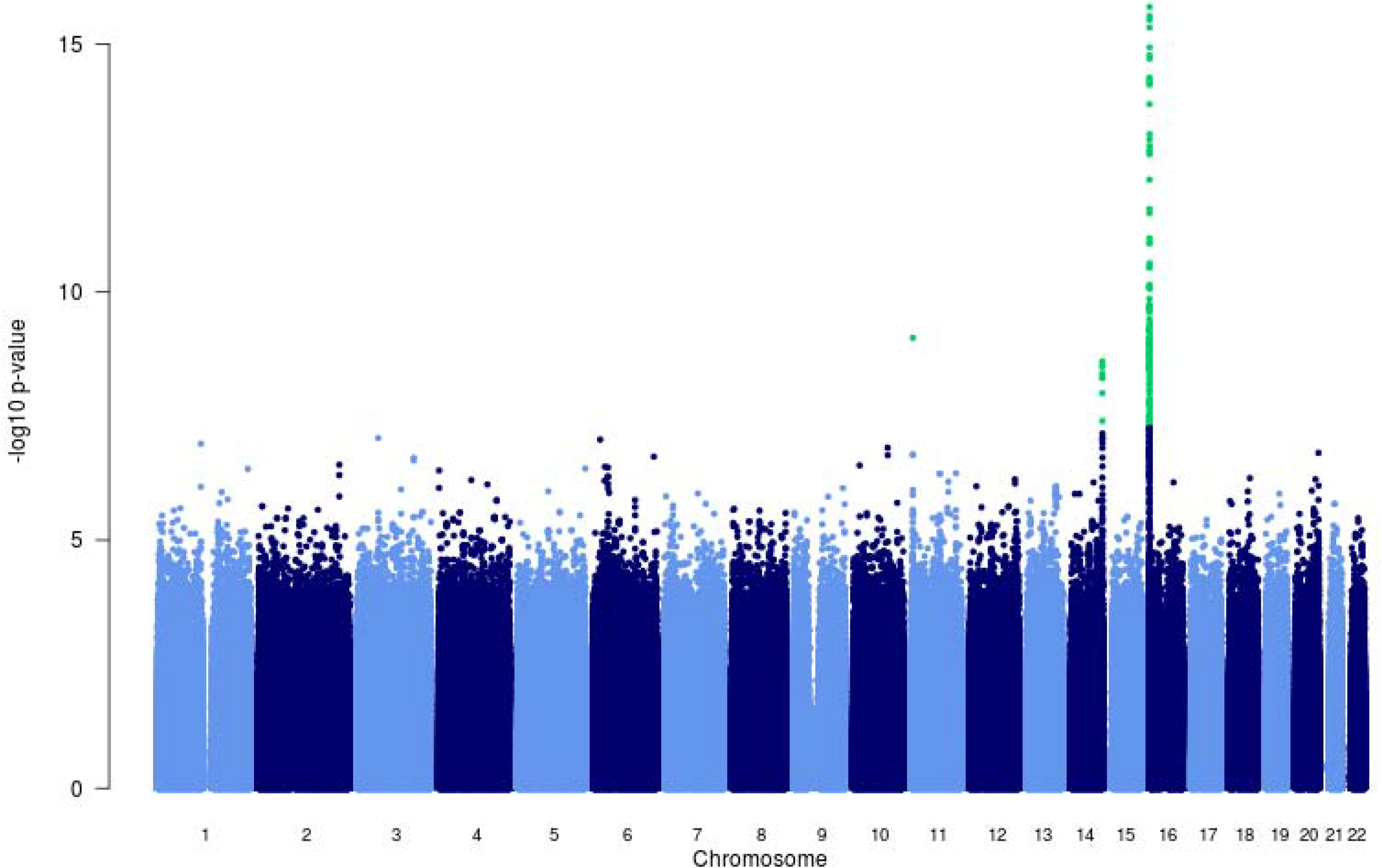
African ancestry hemoglobin analysis Manhattan plot.

**Fig S4.**
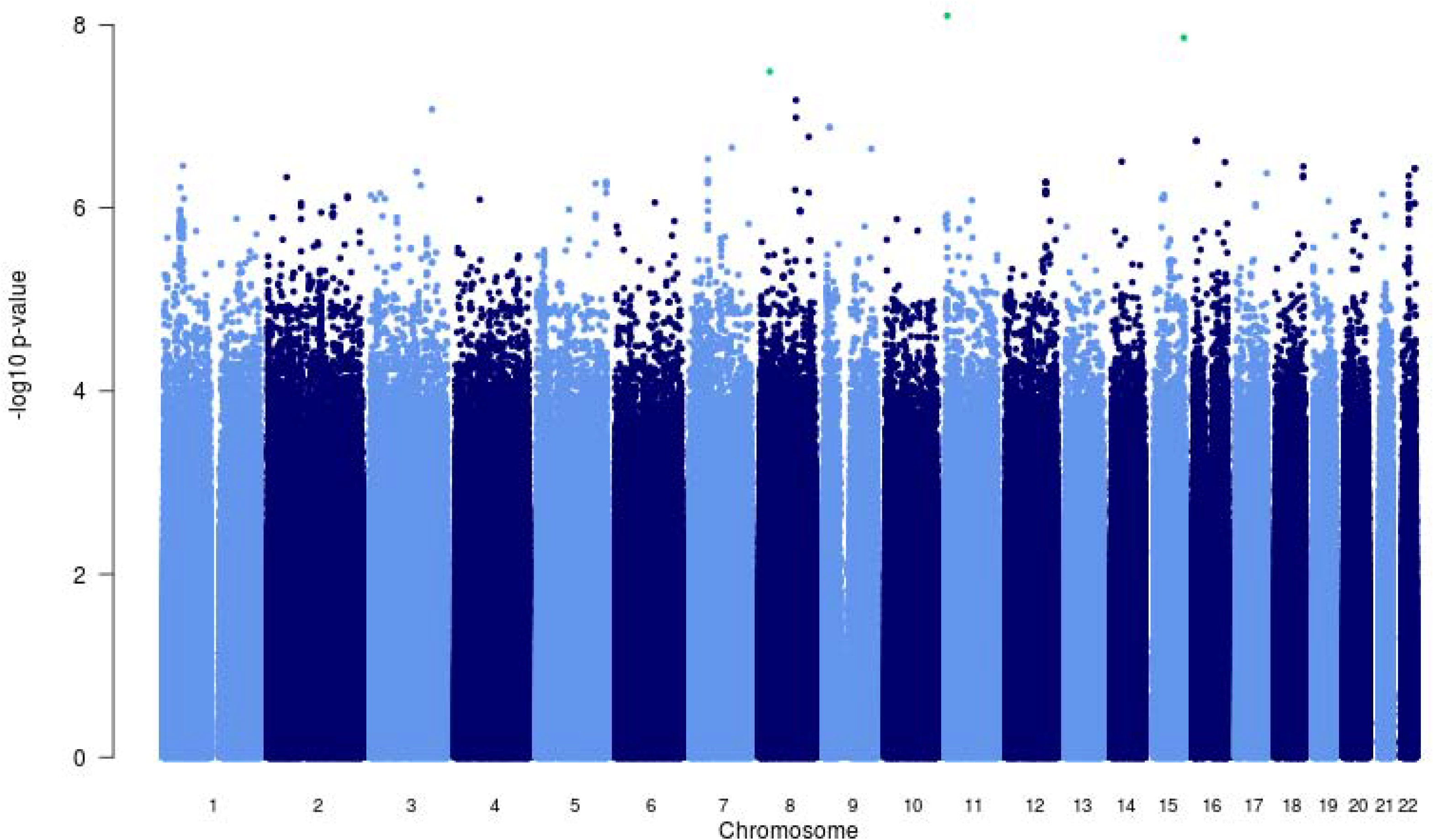
African ancestry white blood cell count analysis Manhattan plot.

**Fig S5.**
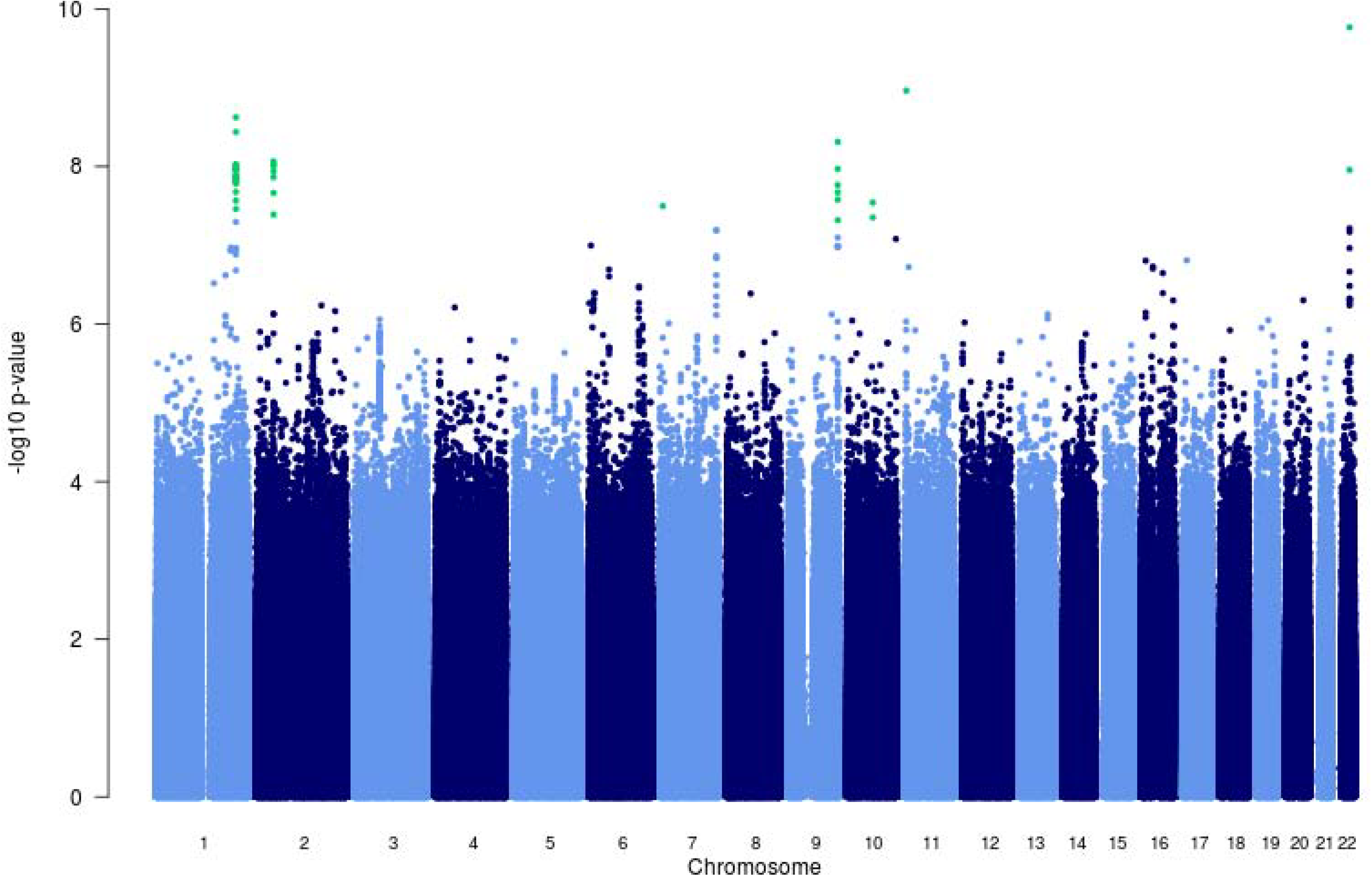
Hispanic/Latino ancestry hematocrit analysis Manhattan plot.

**Fig S6.**
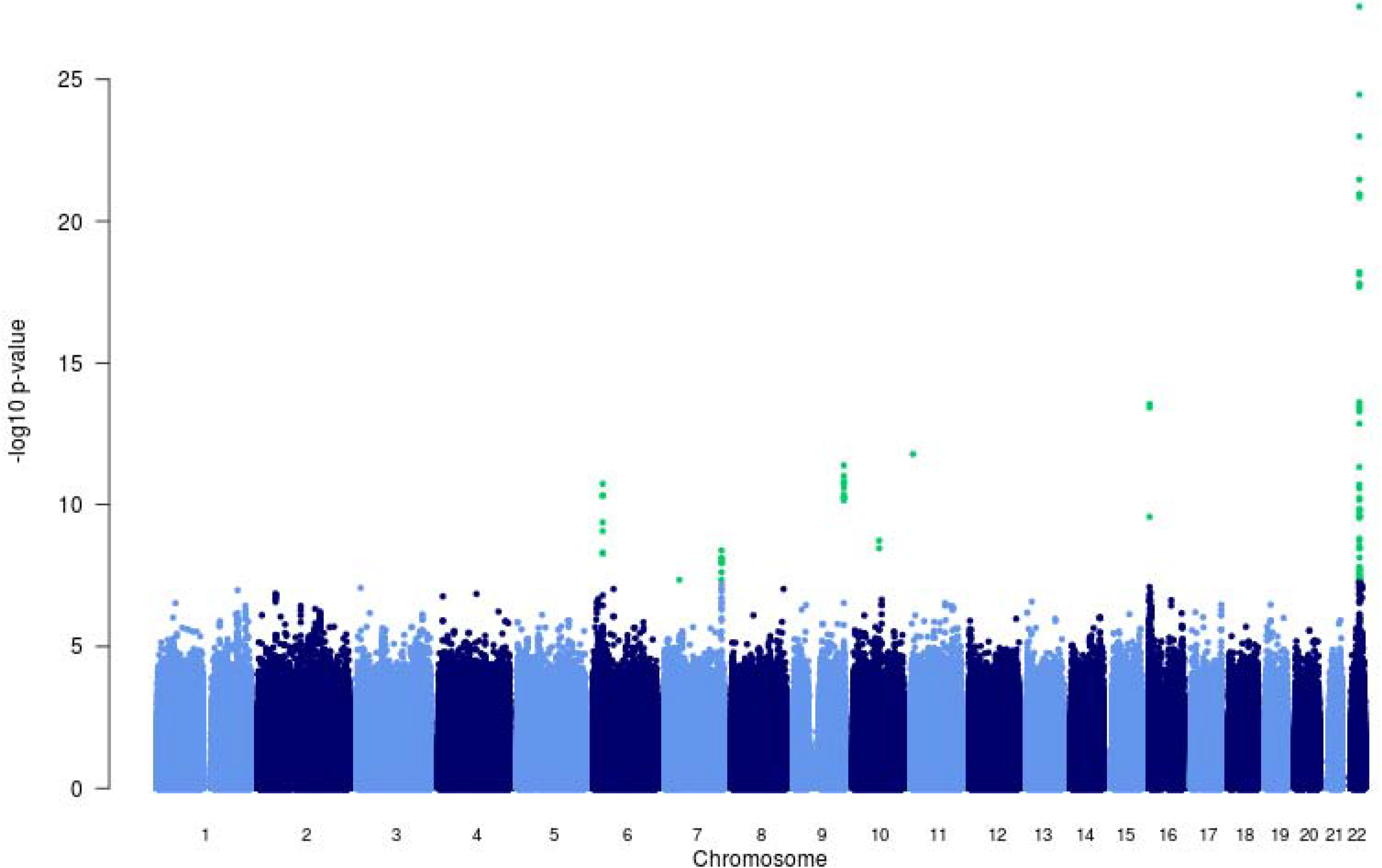
Hispanic/Latino ancestry hemoglobin analysis Manhattan plot.

**Fig S7.**
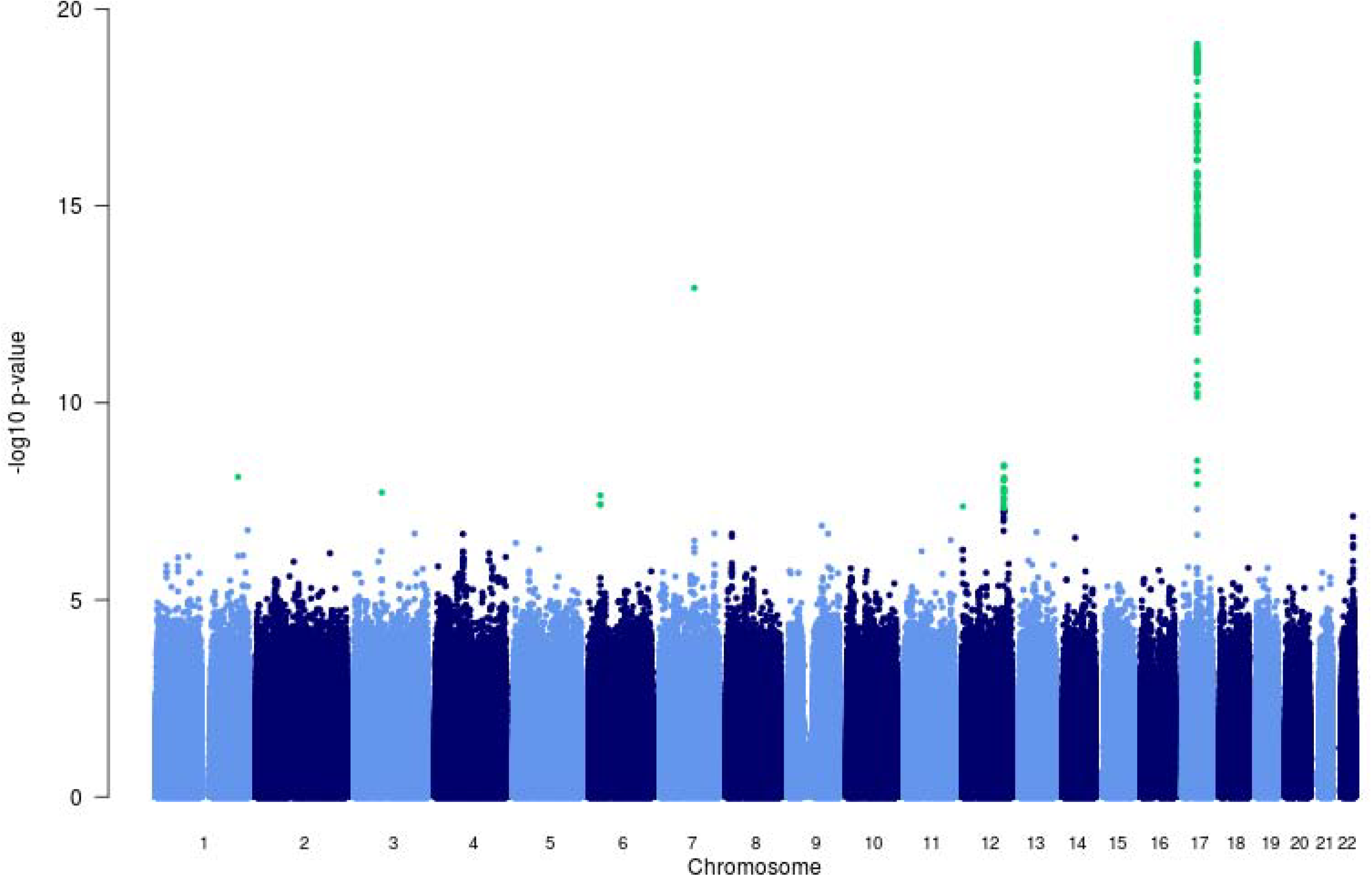
Hispanic/Latino ancestry white blood cell count analysis Manhattan plot.

**Fig S8.**
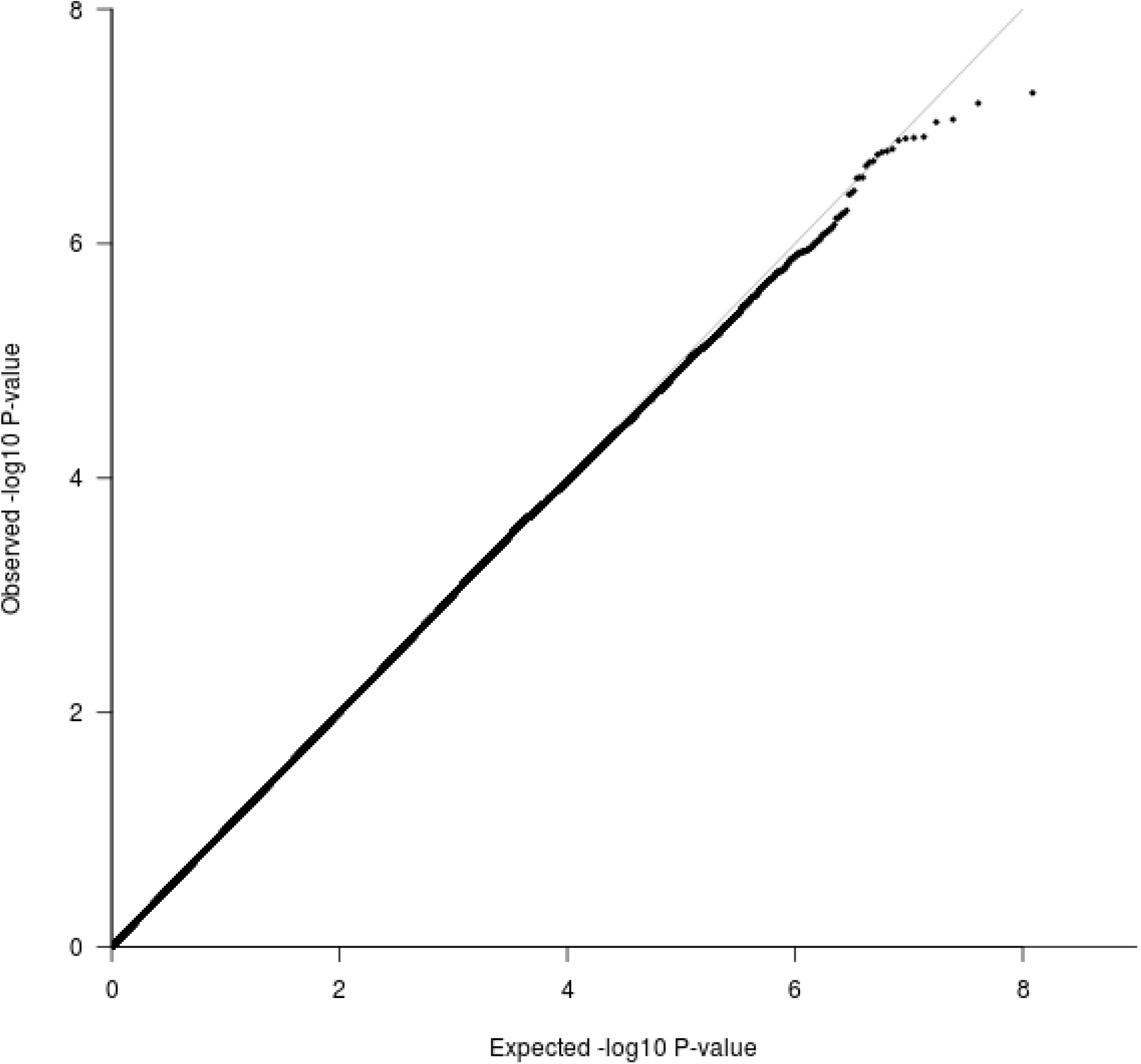
African ancestry hematocrit analysis QQ plot.

**Fig S9.**
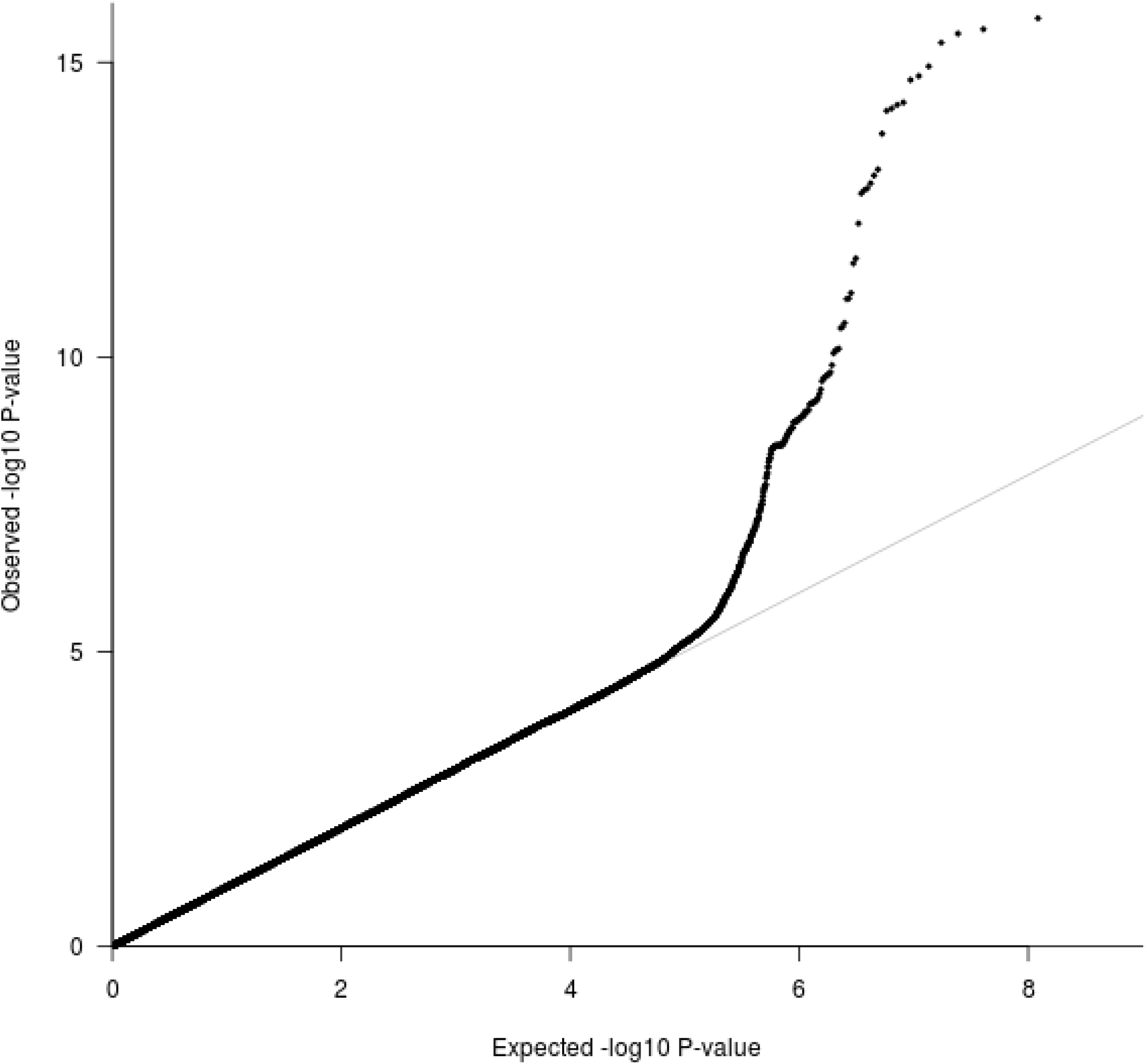
African ancestry hemoglobin analysis QQ plot.

**Fig S10.**
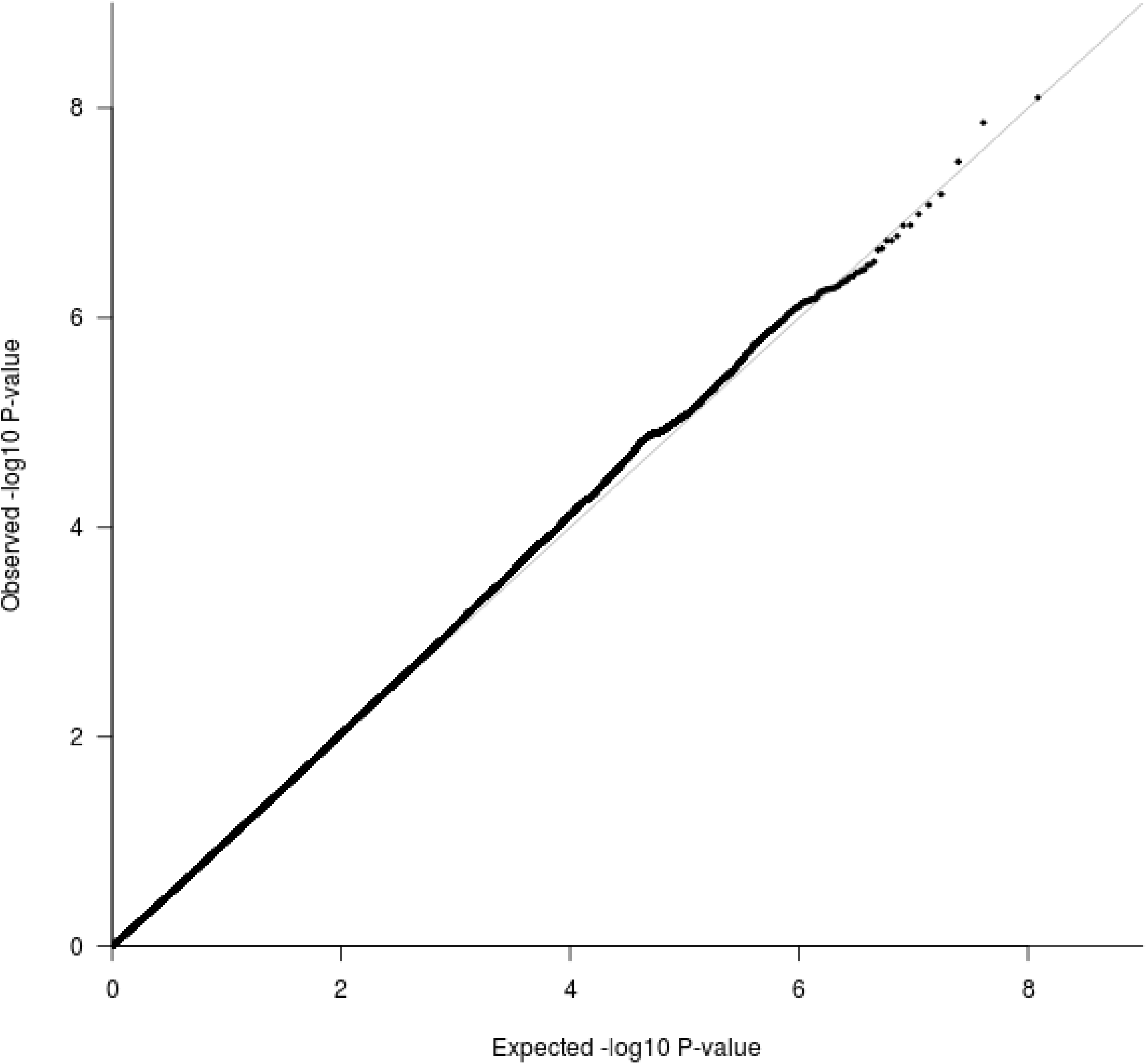
African ancestry white blood cell count analysis QQ plot.

**Fig S11.**
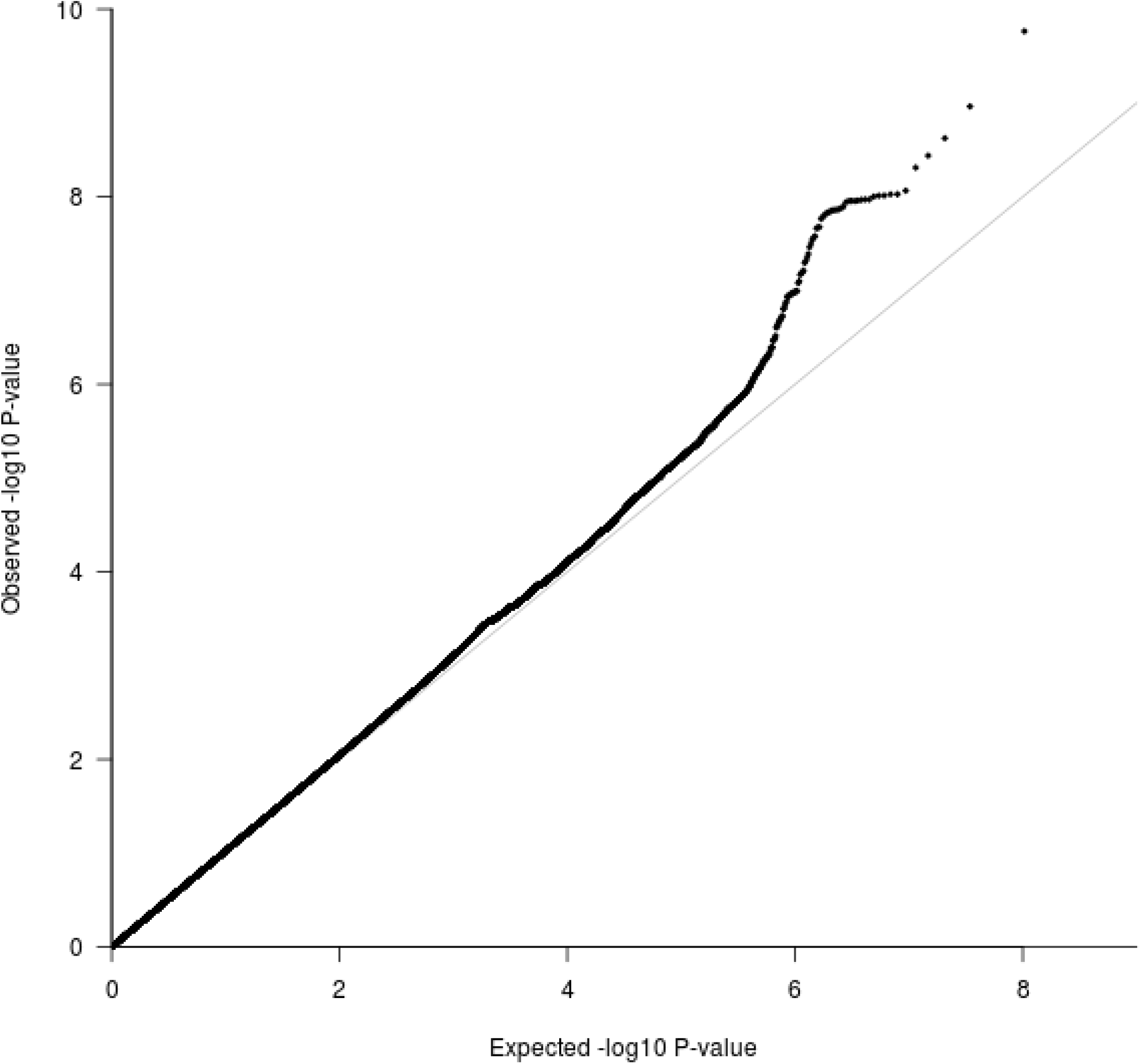
Hispanic/Latino ancestry hematocrit analysis QQ plot.

**Fig S12.**
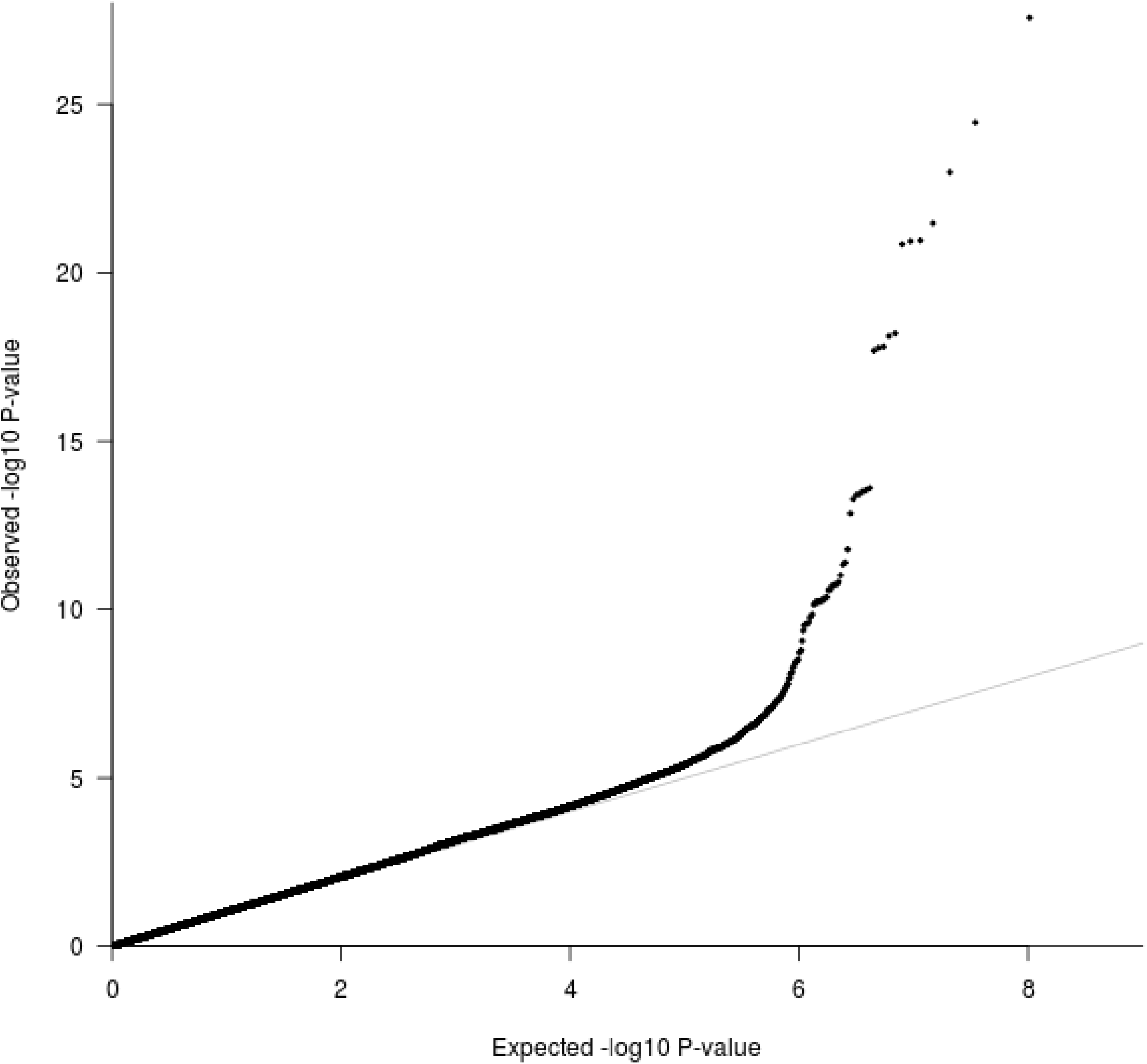
Hispanic/Latino ancestry hemoglobin analysis QQ plot.

**Fig S13.**
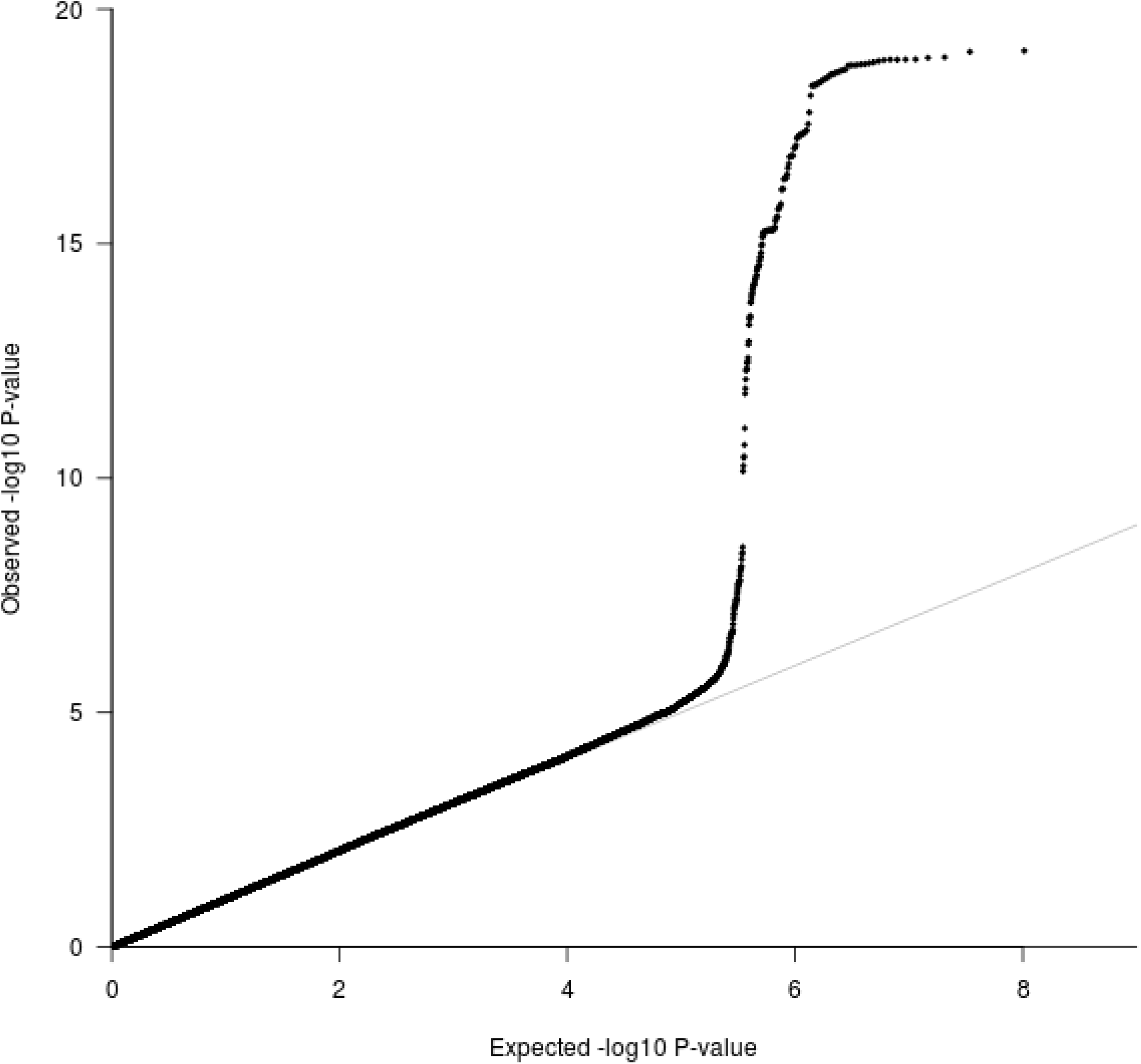
Hispanic/Latino ancestry white blood cell count analysis QQ plot.

**Table S1. Cohorts used for imputation to TOPMed freeze 5b reference panel and subsequent association analysis with hematological traits, including self-identified African ancestry and Hispanic/Latino individuals.**

**Table S2. Cohorts included in the TOPMed freeze 5b imputation reference panels, with self-reported ancestry.**

**Table S3. Imputation quality for variants with a minor allele count between 11 and 20 in Jackson Heart Study.**

**Table S4. Imputation quality for overall reference panel rare variants (MAC<=20 in TOPMed freeze 5b) in Jackson Heart Study.**

**Table S5. Imputation quality for rare variants (MAC<=20) in Hispanic Community Health Study/Study of Latinos (HCHS/SOL).**

**Table S6. Imputation quality for rare variants (MAC<=20) in TOPMed freeze 5b in Hispanic Community Health Study/Study of Latinos (HCHS/SOL).**

**Table S7: Percentage and number of variants well-imputed with TOPMed freeze5b by chromosome in Jackson Heart Study (JHS) and Hispanic Community Health Study/Study of Latinos (HCHS/SOL).**

**Table S8. Demographics, hematological traits, and number of ancestry principal components adjusted for in association analysis models for cohorts imputed to TOPMed freeze 5b reference panel.**

**Table S9. Demographics, hematological traits, and number of ancestry principal components adjusted for in association analysis for African American cohorts with sequencing and hematological trait data from TOPMed freeze 5b.**

**Table S10. Overall counts for variants replicated in our TOPMed freeze 5b imputed cohorts.**

**Table S11. Results for previously identified variants in African ancestry and Hispanic/Latino populations in TOPMed freeze 5b imputed samples (included cohorts detailed in Tables S1 and S8).**

**Table S12: Imputation of novel variants identified with TOPMed freeze 5b-based imputation using current widely used reference panels from the Haplotype Reference Consortium (HRC) and 1000 Genomes Phase 3, as well as subsequent association analysis results for cohorts where the variants were well-imputed (R2>0.8).**

**Table S13. Estimated imputation quality for rs33930165 and rs11549407 using 1000G phase 3 and Haplotype Reference Consortium (HRC) as references.**

**Table S14. Association statistics for the hemoglobin C variant (rs33930165, 11:5227003:C:T) with white blood cell subtypes, adjusting for age, sex, and ancestry principal components.**

**Table S15. White blood cell subtypes for cohorts imputed to TOPMed freeze 5b reference panel.**

**Table S16. White blood cell subtypes for African American cohorts with sequencing and hematological trait data from TOPMed freeze 5b.**

## Supplemental References

1. Auer PL, Johnsen JM, Johnson AD, Logsdon BA, Lange LA, Nalls MA, et al. Imputation of exome sequence variants into population-based samples and blood-cell-trait-associated loci in African Americans: NHLBI GO Exome Sequencing Project. Am J Hum Genet. 2012;91(5):794–808.

2. Duan Q, Liu EY, Auer PL, Zhang G, Lange EM, Jun G, et al. Imputation of coding variants in African Americans: better performance using data from the exome sequencing project. Bioinformatics. 2013;29(21):2744–9.

3. Lu F-P, Lin K-P, Kuo H-K. Diabetes and the risk of multi-system aging phenotypes: a systematic review and meta-analysis. PLoS ONE. 2009;4(1):e4144.

4. Liu EY, Buyske S, Aragaki AK, Peters U, Boerwinkle E, Carlson C, et al. Genotype Imputation of MetabochipSNPs Using a Study-Specific Reference Panel of ~4,000 Haplotypes in African Americans From the Women’s Health Initiative. Genet Epidemiol. 2012;36(2):107–17.

5. Liu EY, Li M, Wang W, Li Y. MaCH-admix: genotype imputation for admixed populations. Genet Epidemiol. 2013;37(1):25–37.

6. Vergara C, Parker MM, Franco L, Cho MH, Valencia-Duarte AV, Beaty TH, et al. Genotype imputation performance of three reference panels using African ancestry individuals. Hum Genet. 2018;137(4):281–92.

7. The International HapMap Consortium. A second generation human haplotype map of over 3.1 million SNPs. Nature. 2007;449:851–61.

8. McCarthy S, Das S, Kretzschmar W, Delaneau O, Wood AR, Teumer A, et al. A reference panel of 64,976 haplotypes for genotype imputation. Nat Genet. 2016;48(10):1279–83.

9. The 1000 Genomes Project Consortium. A global reference for human genetic variation. Nature. 2015;526(7571):68–74.

10. Mathias RA, Taub MA, Gignoux CR, Fu W, Musharoff S, O’Connor TD, et al. A continuum of admixture in the Western Hemisphere revealed by the African Diaspora genome. Nature communications. 2016;7:12522.

11. Crosslin DR, McDavid A, Weston N, Nelson SC, Zheng X, Hart E, et al. Genetic variants associated with the white blood cell count in 13,923 subjects in the eMERGE Network. Hum Genet. 2012;131(4):639–52.

12. Whitfield JB, Martin NG, Rao DC. Genetic and environmental influences on the size and number of cells in the blood. Genetic epidemiology. 1985;2(2):133–44.

13. Garner C, Tatu T, Reittie JE, Littlewood T, Darley J, Cervino S, et al. Genetic influences on F cells and other hematologic variables: a twin heritability study. Blood. 2000;95(1):342–6.

14. Astle WJ, Elding H, Jiang T, Allen D, Ruklisa D, Mann AL, et al. The Allelic Landscape of Human Blood Cell Trait Variation and Links to Common Complex Disease. Cell. 2016;167(5):1415–29 e19.

15. van der Harst P, Zhang W, Mateo Leach I, Rendon A, Verweij N, Sehmi J, et al. Seventy-five genetic loci influencing the human red blood cell. Nature. 2012;492(7429):369–75.

16. Mousas A, Ntritsos G, Chen MH, Song C, Huffman JE, Tzoulaki I, et al. Rare coding variants pinpoint genes that control human hematological traits. PLoS Genet. 2017;13(8):e1006925.

17. Chami N, Chen MH, Slater AJ, Eicher JD, Evangelou E, Tajuddin SM, et al. Exome Genotyping Identifies Pleiotropic Variants Associated with Red Blood Cell Traits. Am J Hum Genet. 2016.

18. Eicher JD, Chami N, Kacprowski T, Nomura A, Chen MH, Yanek LR, et al. Platelet-Related Variants Identified by Exomechip Meta-analysis in 157,293 Individuals. Am J Hum Genet. 2016.

19. Tajuddin SM, Schick UM, Eicher JD, Chami N, Giri A, Brody JA, et al. Large-Scale Exome-wide Association Analysis Identifies Loci for White Blood Cell Traits and Pleiotropy with Immune-Mediated Diseases. Am J Hum Genet. 2016.

20. Hodonsky CJ, Jain D, Schick UM, Morrison JV, Brown L, McHugh CP, et al. Genome-wide association study of red blood cell traits in Hispanics/Latinos: The Hispanic Community Health Study/Study of Latinos. PLoS Genet. 2017;13(4):e1006760.

21. Group. CCHW. Meta-analysis of rare and common exome chip variants identifies S1PR4 and other loci influencing blood cell traits. Nat Genet. 2016;48(8):867–76.

22. Lo KS, Wilson JG, Lange LA, Folsom AR, Galarneau G, Ganesh SK, et al. Genetic association analysis highlights new loci that modulate hematological trait variation in Caucasians and African Americans. Hum Genet. 2011;129(3):307–17.

23. Tournamille C, Colin Y, Cartron JP, Le Van Kim C. Disruption of a GATA motif in the Duffy gene promoter abolishes erythroid gene expression in Duffy-negative individuals. Nat Genet. 1995;10(2):224–8.

24. Chen Z, Tang H, Qayyum R, Schick UM, Nalls MA, Handsaker R, et al. Genome-wide association analysis of red blood cell traits in African Americans: the COGENT Network. Hum Mol Genet. 2013;22(12):2529–38.

25. Li J, Glessner JT, Zhang H, Hou C, Wei Z, Bradfield JP, et al. GWAS of blood cell traits identifies novel associated loci and epistatic interactions in Caucasian and African-American children. Hum Mol Genet. 2013;22(7):1457–64.

26. van Rooij FJA, Qayyum R, Smith AV, Zhou Y, Trompet S, Tanaka T, et al. Genome-wide Trans-ethnic Meta-analysis Identifies Seven Genetic Loci Influencing Erythrocyte Traits and a Role for RBPMS in Erythropoiesis. Am J Hum Genet. 2017;100(1):51–63.

27. Jain D, Hodonsky CJ, Schick UM, Morrison JV, Minnerath S, Brown L, et al. Genome-wide association of white blood cell counts in Hispanic/Latino Americans: the Hispanic Community Health Study/Study of Latinos. Hum Mol Genet. 2017;26(6):1193–204.

28. Polfus LM, Khajuria RK, Schick UM, Pankratz N, Pazoki R, Brody JA, et al. Whole-Exome Sequencing Identifies Loci Associated with Blood Cell Traits and Reveals a Role for Alternative GFI1B Splice Variants in Human Hematopoiesis. Am J Hum Genet. 2016;99(2):481–8.

29. Keller MF, Reiner AP, Okada Y, van Rooij FJ, Johnson AD, Chen MH, et al. Trans-ethnic meta-analysis of white blood cell phenotypes. Hum Mol Genet. 2014;23(25):6944–60.

30. Reiner AP, Lettre G, Nalls MA, Ganesh SK, Mathias R, Austin MA, et al. Genome-wide association study of white blood cell count in 16,388 African Americans: the continental origins and genetic epidemiology network (COGENT). PLoS Genet. 2011;7(6):e1002108.

31. Pulit SL, de With SA, de Bakker PI. Resetting the bar: Statistical significance in whole-genome sequencing-based association studies of global populations. Genet Epidemiol. 2017;41(2):145–51.

32. Trecartin RF, Liebhaber SA, Chang JC, Lee KY, Kan YW, Furbetta M, et al. beta zero thalassemia in Sardinia is caused by a nonsense mutation. The Journal of clinical investigation. 1981;68(4):1012–7.

33. Rosatelli MC, Dozy A, Faa V, Meloni A, Sardu R, Saba L, et al. Molecular characterization of beta-thalassemia in the Sardinian population. Am J Hum Genet. 1992;50(2):422–6.

34. Perea FJ, Magana MT, Cobian JG, Sanchez-Lopez JY, Chavez ML, Zamudio G, et al. Molecular spectrum of beta-thalassemia in the Mexican population. Blood cells, molecules & diseases. 2004;33(2):150–2.

35. Silva AN, Cardoso GL, Cunha DA, Diniz IG, Santos SE, Andrade GB, et al. The Spectrum of beta-Thalassemia Mutations in a Population from the Brazilian Amazon. Hemoglobin. 2016;40(1):20–4.

36. Key NS, Connes P, Derebail VK. Negative health implications of sickle cell trait in high income countries: from the football field to the laboratory. British journal of haematology. 2015;170(1):5–14.

37. Graffeo L, Vitrano A, Scondotto S, Dardanoni G, Pollina Addario WS, Giambona A, et al. beta-Thalassemia heterozygote state detrimentally affects health expectation. European journal of internal medicine. 2018;54:76–80.

38. Galanello R, Origa R. Beta-thalassemia. Orphanet journal of rare diseases. 2010;5:11-.

39. Fairhurst RM, Casella JF. Images in clinical medicine. Homozygous hemoglobin C disease. Q1‘‘. 2004;350(26):e24.

40. Sorlie PD, Aviles-Santa LM, Wassertheil-Smoller S, Kaplan RC, Daviglus ML, Giachello AL, et al. Design and implementation of the Hispanic Community Health Study/Study of Latinos. Ann Epidemiol. 2010;20(8):629–41.

41. Daviglus ML, Talavera GA, Aviles-Santa ML, Allison M, Cai J, Criqui MH, et al. Prevalence of major cardiovascular risk factors and cardiovascular diseases among Hispanic/Latino individuals of diverse backgrounds in the United States. JAMA. 2012;308(17):1775–84.

42. Lavange LM, Kalsbeek WD, Sorlie PD, Aviles-Santa LM, Kaplan RC, Barnhart J, et al. Sample design and cohort selection in the Hispanic Community Health Study/Study of Latinos. Ann Epidemiol. 2010;20(8):642–9.

43. Conomos MP, Laurie CA, Stilp AM, Gogarten SM, McHugh CP, Nelson SC, et al. Genetic Diversity and Association Studies in US Hispanic/Latino Populations: Applications in the Hispanic Community Health Study/Study of Latinos. Am J Hum Genet. 2016;98(1):165–84.

44. Wojcik G, Graff M, Nishimura KK, Tao R, Haessler J, Gignoux CR, et al. The PAGE Study: How Genetic Diversity Improves Our Understanding of the Architecture of Complex Traits. bioRxiv. 2018:188094.

45. Design of the Women’s Health Initiative clinical trial and observational study. The Women’s Health Initiative Study Group. Control Clin Trials. 1998;19(1):61–109.

46. UK Biobank. UK Biobank: rationale, design and development of a large-scale prospective resource. 2007. [Available from: http://www.ukbiobank.ac.uk/resources/.

47. Bycroft C, Freeman C, Petkova D, Band G, Elliott LT, Sharp K, et al. The UK Biobank resource with deep phenotyping and genomic data. Nature. 2018;562(7726):203–9.

48. Kvale MN, Hesselson S, Hoffmann TJ, Cao Y, Chan D, Connell S, et al. Genotyping Informatics and Quality Control for 100,000 Subjects in the Genetic Epidemiology Research on Adult Health and Aging (GERA) Cohort. Genetics. 2015;200(4):1051–60.

49. Banda Y, Kvale MN, Hoffmann TJ, Hesselson SE, Ranatunga D, Tang H, et al. Characterizing Race/Ethnicity and Genetic Ancestry for 100,000 Subjects in the Genetic Epidemiology Research on Adult Health and Aging (GERA) Cohort. Genetics. 2015;200(4):1285–95.

50. Taylor HA, Jr., Wilson JG, Jones DW, Sarpong DF, Srinivasan A, Garrison RJ, et al. Toward resolution of cardiovascular health disparities in African Americans: design and methods of the Jackson Heart Study. Ethn Dis. 2005;15(4 Suppl 6):S6–4-17.

51. Wilson JG, Rotimi CN, Ekunwe L, Royal CD, Crump ME, Wyatt SB, et al. Study design for genetic analysis in the Jackson Heart Study. Ethn Dis. 2005;15(4 Suppl 6):S6–30-7.

52. Musunuru K, Lettre G, Young T, Farlow DN, Pirruccello JP, Ejebe KG, et al. Candidate gene association resource (CARe): design, methods, and proof of concept. Circulation Cardiovascular genetics. 2010;3(3):267–75.

53. Lettre G, Palmer CD, Young T, Ejebe KG, Allayee H, Benjamin EJ, et al. Genome-wide association study of coronary heart disease and its risk factors in 8,090 African Americans: the NHLBI CARe Project. PLoS Genet. 2011;7(2):e1001300.

54. Friedman GD, Cutter GR, Donahue RP, Hughes GH, Hulley SB, Jacobs DR, Jr., et al. CARDIA: study design, recruitment, and some characteristics of the examined subjects. J Clin Epidemiol. 1988;41(11):1105–16.

55. Cutter GR, Burke GL, Dyer AR, Friedman GD, Hilner JE, Hughes GH, et al. Cardiovascular risk factors in young adults. The CARDIA baseline monograph. Control Clin Trials. 1991;12(1 Suppl):1S–77S.

56. The Atherosclerosis Risk in Communities (ARIC) Study: design and objectives. The ARIC investigators. Am J Epidemiol. 1989;129(4):687–702.

57. Loh PR, Danecek P, Palamara PF, Fuchsberger C, Y Ar, H Kf, et al. Reference-based phasing using the Haplotype Reference Consortium panel. Nat Genet. 2016;48(11):1443–8.

58. Das S, Forer L, Schonherr S, Sidore C, Locke AE, Kwong A, et al. Next-generation genotype imputation service and methods. Nat Genet. 2016;48(10):1284–7.

59. Duan Q, Liu EY, Croteau-Chonka DC, Mohlke KL, Li Y. A comprehensive SNP and indel imputability database. Bioinformatics. 2013;29(4):528–31.

60. Magi R, Lindgren CM, Morris AP. Meta-analysis of sex-specific genome-wide association studies. Genet Epidemiol. 2010;34(8):846–53.

61. Maples Brian K, Gravel S, Kenny Eimear E, Bustamante Carlos D. RFMix: A Discriminative Modeling Approach for Rapid and Robust Local-Ancestry Inference. The American Journal of Human Genetics. 2013;93(2):278–88.

62. Li JZ, Absher DM, Tang H, Southwick AM, Casto AM, Ramachandran S, et al. Worldwide human relationships inferred from genome-wide patterns of variation. Science. 2008;319(5866):1100–4.

